# Bioisosteric analogs of MDMA with improved pharmacological profile

**DOI:** 10.1101/2024.04.08.588083

**Authors:** Ana Sofia Alberto-Silva, Selina Hemmer, Hailey A. Bock, Leticia Alves da Silva, Kenneth R. Scott, Nina Kastner, Manan Bhatt, Marco Niello, Kathrin Jäntsch, Oliver Kudlacek, Elena Bossi, Thomas Stockner, Markus R. Meyer, John D. McCorvy, Simon D. Brandt, Pierce Kavanagh, Harald H. Sitte

## Abstract

3,4-Methylenedioxymethamphetamine (MDMA, ‘*ecstasy’*) is re-emerging in clinical settings as a candidate for the treatment of specific psychiatric disorders (e.g. post-traumatic stress disorder) in combination with psychotherapy. MDMA is a psychoactive drug, typically regarded as an empathogen or entactogen, which leads to transporter-mediated monoamine release. Despite its therapeutic potential, MDMA can induce dose-, individual-, and context-dependent untoward effects outside safe settings. In this study, we investigated whether three new methylenedioxy bioisosteres of MDMA improve its off-target profile. *In vitro* methods included radiotracer assays, transporter electrophysiology, bioluminescence resonance energy transfer and fluorescence-based assays, pooled human liver microsome/S9 fraction incubation with isozyme mapping, and liquid chromatography coupled to high-resolution mass spectrometry. *In silico* methods included molecular docking. Compared with MDMA, all three MDMA bioisosteres (ODMA, TDMA, and SeDMA) showed similar pharmacological activity at human serotonin and dopamine transporters (hSERT and hDAT, respectively) but decreased activity at 5-HT_2A/2B/2C_ receptors. Regarding their hepatic metabolism, they differed from MDMA, with *N*-demethylation being the only metabolic route shared, and without forming phase II metabolites. Additional screening for their interaction with human organic cation transporters (hOCTs) and plasma membrane transporter (hPMAT) revealed a weaker interaction of the MDMA analogs with hOCT1, hOCT2, and hPMAT. Our findings suggest that these new MDMA analogs might constitute appealing therapeutic alternatives to MDMA, sparing the primary pharmacological activity at hSERT and hDAT, but displaying a reduced activity at 5-HT_2A/2B/2C_ receptors and reduced hepatic metabolism. Whether these MDMA bioisosteres may pose lower risk alternatives to the clinically re-emerging MDMA warrants further studies.

## 1. Introduction

3,4-Methylenedioxymethamphetamine (MDMA, also known as ‘*ecstasy’*) (Fig. 1A) is a psychoactive drug capable of inducing a “controlled altered state of consciousness” (Shulgin and Nichols 1978). In recent years, MDMA re-emerged in preclinical and clinical research for the treatment of specific neuropsychiatric disorders, such as post-traumatic stress disorder (PTSD), in combination with psychotherapy (Mithoefer *et al*. 2011; Mithoefer *et al*. 2018; Mitchell *et al*. 2021; Mitchell *et al*. 2023).

**Fig. 1.**
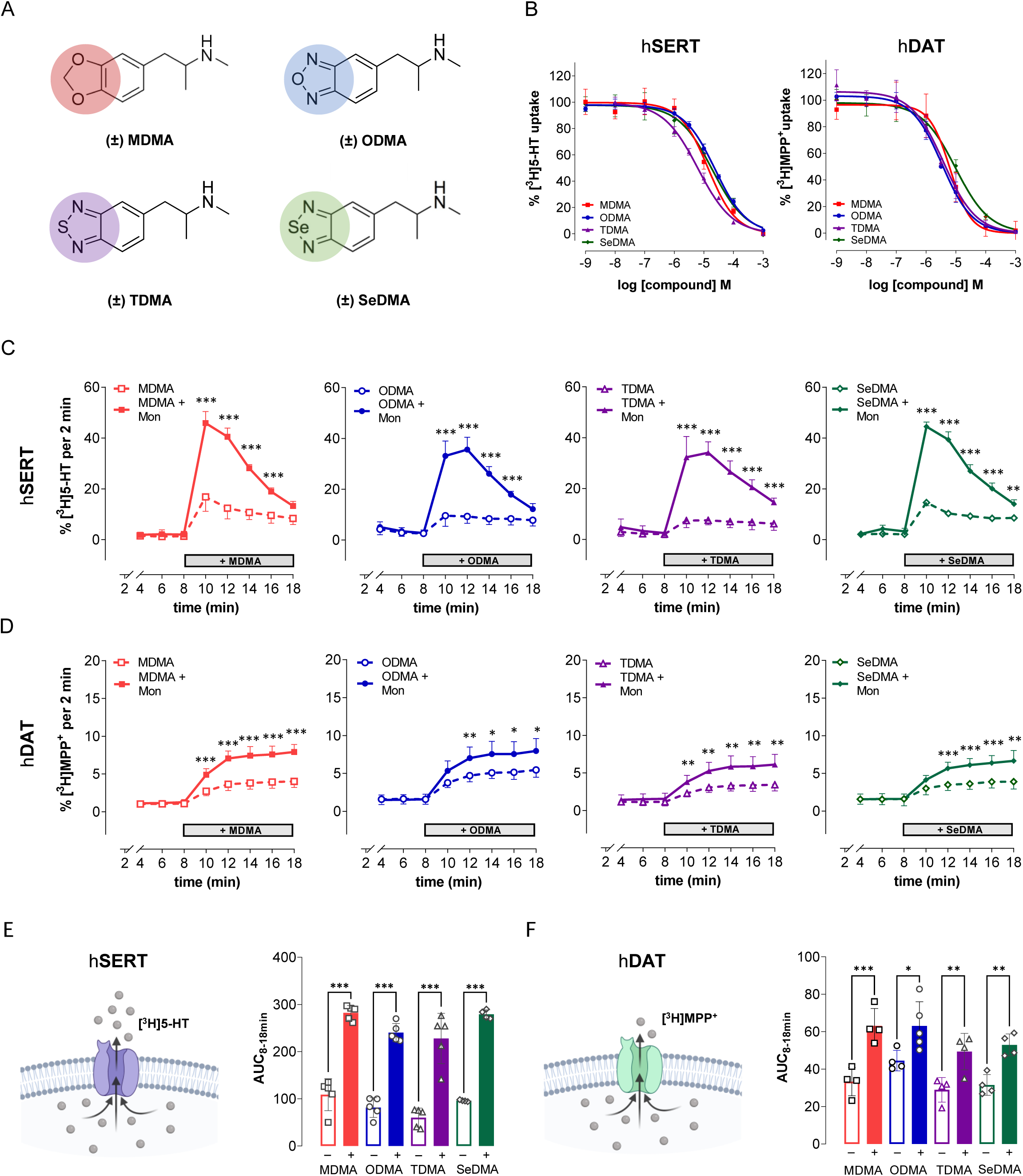
MDMA and its analogs ODMA, TDMA, and SeDMA interact at low micromolar concentrations with the human serotonin and dopamine transporters. **(A)** Chemical structures of MDMA and its analogs ODMA, TDMA, and SeDMA, in which the methylenedioxy group or its chemical modification is highlighted in red, blue, purple, or green, respectively; **(B)** Uptake inhibition curves at hSERT (left panel) and hDAT (right panel). Curves were plotted and fitted by non-linear regression, and data were best fitted to a sigmoidal dose-response curve to obtain IC_50_ values (Table 1); **(C)** Transporter-mediated release (batch) of preloaded [^3^H]5-HT from HEK293 cells stably expressing hSERT. Compounds were added at a concentration close to their IC_50_ value (MDMA (14 µM), ODMA (23 µM), TDMA (7 µM), SeDMA (20 µM)) from 8 to 18 min, either in KHB (empty symbols) or in KHB+Mon (10 μM) (filled symbols); **(D)** Transporter-mediated release (superfusion) of preloaded [^3^H]MPP^+^ from HEK293 cells stably expressing hDAT. Similarly, compounds were added at a concentration close to their IC_50_ value (MDMA, ODMA, and TDMA (6 µM), SeDMA (10 µM)) from 8 to 18 min either in KHB (empty symbols) or in KHB+Mon (10 μM) (filled symbols); **(E)** Illustrative representation of [^3^H]5-HT reverse transport at hSERT (left panel; created with BioRender.com) and the calculated area under the curve between 8 and 18 minutes (AUC_8-18min_) of the [^3^H]5-HT released by each compound in KHB (-) or KHB+Mon (+) conditions (right panel) (MDMA_KHB_ = 108.9 ± 34.0 vs. MDMA_MON_ = 282.3 ± 15.4; ODMA_KHB_ = 81.5 ± 20.9 vs. ODMA_MON_ = 240.3 ± 19.3; TDMA_KHB_ = 60.2 ± 20.0 vs. TDMA_MON_ = 227.9 ± 53.4; SeDMA_KHB_ = 96.0 ± 1.7 vs. SeDMA_MON_ = 279.2 ± 9.3); (**F**) Illustrative representation of [^3^H]MPP^+^ reverse transport at hDAT (left panel; created with BioRender.com) and the calculated AUC_8-18min_ of the [^3^H]MPP^+^ released by each compound in KHB (-) or KHB+Mon (+) conditions (right panel) (MDMA_KHB_ = 33.3 ± 7.5 vs. MDMA_MON_ = 63.1 ± 9.3; ODMA_KHB_ = 44.5 ± 5.5 vs. ODMA_MON_ = 63.1 ± 13.0; TDMA_KHB_ = 28.9 ± .6.6 vs. TDMA_MON_ = 49.4 ± 9.7; SeDMA_KHB_ = 31.6 ± 5.5 vs. SeDMA_MON_ = 52.9 ± 5.9). Data are mean ± SD from three to five individual experiments, performed in triplicate (uptake inhibition assays and superfusion release assays) or duplicate (batch release assays). Statistical analyses explored possible significant differences between KHB and KHB+Mon conditions. Statistical significance was defined at a p-value lower than 0.05. *denotes p<0.05, **p<0.01, and ***p<0.001.

**Table 1.**
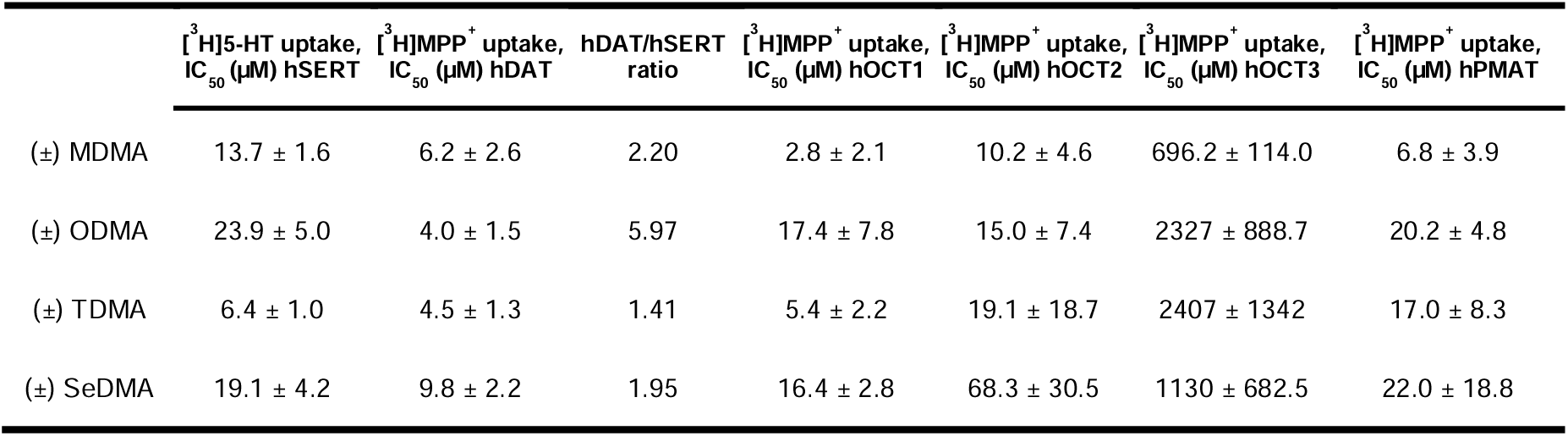
Uptake inhibition assays. Potency of MDMA and its analogs ODMA, TDMA, and SeDMA at hSERT, hDAT, hOCT1, hOCT2, hOCT3, and hPMAT. Data represent mean and SD from at least three independent experiments performed in triplicate. DAT/SERT ratio = (1/DAT_IC50_):(1/SERT_IC50_).

MDMA is a ring-substituted amphetamine derivative with psychostimulant activity. MDMA is unique in inducing an interoceptive and prosocial effect, and it has been described as an empathogen or entactogen (Nichols 1986). Although its mechanism of action is not yet fully elucidated, MDMA is generally recognized to interact with monoamine transporters for serotonin (SERT), dopamine (DAT), and norepinephrine (NET), eliciting non-exocytotic efflux of serotonin (5-HT), dopamine (DA) and norepinephrine (NE), respectively (Rudnick and Wall 1992; Rothman *et al*. 2001). Additionally, MDMA is an agonist at 5-HT_2A/2B/2C_ receptors (Nash *et al*. 1994; Setola *et al*. 2003). The reported acute and chronic side effects of MDMA can range from tachycardia and hypertension to hyperthermia, cardiotoxicity, and hepatotoxicity (Vollenweider *et al*. 1998; Delaforge *et al*. 1999; Setola *et al*. 2003; De La Torre *et al*. 2004; Capela *et al*. 2006b; Capela *et al*. 2006a; Bhattacharyya *et al*. 2009; Steinkellner *et al*. 2011; Vizeli *et al*. 2017; Dunlap *et al*. 2018; Fonseca *et al*. 2021). MDMA is rapidly absorbed in the intestinal tract and its metabolism displays non-linear pharmacokinetics, which has been partially linked to the inhibition of certain cytochrome P450 (CYP) enzymes (De La Torre *et al*. 2004; Yang *et al*. 2006; Dunlap *et al*. 2018). This enzymatic inhibition has mainly been associated with the methylenedioxy group of MDMA (Delaforge *et al*. 1999; De La Torre *et al*. 2004; Dinger *et al*. 2016). Additionally, the metabolites of MDMA have been described to be responsible for MDMA-related neurotoxicity in rodents, since bypassing metabolism through direct intracerebroventricular administration of MDMA did not induce neurotoxicity (Paris and Cunningham 1992; Esteban *et al*. 2001; Green *et al*. 2003). Moreover, it was reported that MDMA metabolites (mainly catechol and quinone metabolites formed after opening of the methylenedioxy group) could generate free radicals (e.g. reactive oxygen species), which might induce oxidative stress and cellular damage (Jayanthi *et al*. 1999; Shankaran *et al*. 1999; Capela *et al*. 2006a).

In this study, we investigated three new MDMA analogs with a bioisosteric replacement of the methylenedioxyphenyl (or 1,3-benzodioxole) group of MDMA. This chemical modification has been described to be able to evade the inhibition of CYP enzymes, namely CYP2D6 (Meanwell 2014; Anzali *et al*. 1997). The analogs were designed by the replacement of the 1,3-benzodioxole group with 2,1,3-benzoxadiazole, 2,1,3-benzothiadiazole, and 2,1,3-benzoselenadiazole, which gave 1-(2,1,3-benzoxadiazol-5-yl)-*N*-methylpropan-2-amine (ODMA), 1-(2,1,3-benzothiadiazol-5-yl)-*N*-methylpropan-2-amine (TDMA), and 1-(2,1,3-benzoselenadiazol-5-yl)-*N*-methylpropan-2-amine (SeDMA), respectively (Fig. 1A). The main aims of this study were to characterize the molecular mode of action of these three analogs at key targets: monoamine transporters (specifically SERT and DAT), a subset of serotonin receptors (subfamily 2), organic cation transporters, and plasma membrane transporters. In addition, we studied the *in vitro* hepatic metabolism of these MDMA analogs and how it differed from MDMA.

Considering the reported MDMA-induced adverse events, it is advantageous to explore MDMA-related congeners which can putatively keep or improve its therapeutic action but potentially decrease its off-target effects.

## 2. Materials and methods

See the **Supplementary Information** for the complete details in each section.

### 2.1. Drugs and reagents

The experimental drug MDMA hydrochloride (HCl) (MW = 229.7 g/mol) was purchased from Lipomed AG (Arlesheim, Switzerland) or Cayman Chemical (Ann Arbor, MI, USA). ODMA HCl (MW = 227.69 g/mol), TDMA HCl (MW = 243.75 g/mol) and SeDMA succinate (MW = 254.19:118.09 g/mol) (≥95%) were synthesized using established methods (Abdel-Magid *et al*. 1996; Briner *et al*. 2000; Gadakh *et al*. 2014). Identity and purities were confirmed by standard analytical characterizations. All MDMA and analogs were racemates (±; *R/S*). Other experimental drugs comprised paroxetine hydrochloride (HCl), vanoxerine (GBR12909), *para*-chloroamphetamine (*p*CA) HCl, dextroamphetamine hemisulfate salt (d-amp; *(S)*-amphetamine), monensin (Mon) and dopamine (DA), and were supplied by Sigma-Aldrich (St. Louis, MO, United States). Serotonin (5-hydroxytryptamine; 5-HT) HCl was obtained from Fluorochem Ltd (Hadfield, United Kingdom). For cell culture, Dulbecco’s Modified Eagle Medium (DMEM) High Glucose (4.5 g/L) with L-glutamine and fetal bovine serum (FBS) were obtained from Capricorn Scientific GmbH (Ebsdorfergrund, Germany), as well as geneticin (G-418 sulfate solution; 50 mg/mL). Blasticidin (10 mg/mL) and zeocin (100 mg/mL) were purchased from InvivoGen (San Diego, CA, United States). Tetracycline HCl was obtained from former Boehringer Mannheim (Mannheim, Germany). Penicillin-streptomycin (10 000 IU/10 mg/100 mL) was purchased from Sigma-Aldrich (St. Louis, MO, United States). For radiolabeled assays, [^3^H]5-HT and [^3^H]1-methyl-4-phenylpyridinium ([^3^H]MPP^+^) were obtained from PerkinElmer, Inc (Waltham, MA, USA).

### 2.2. Cell culture

Human embryonic kidney 293 (HEK293) cells stably expressing the human isoforms of SERT, DAT, OCT1-3, and PMAT were used. YFP-tagged constructs were used in uptake-inhibition and release assays. The generation and maintenance of stable cell lines expressing SERT or DAT were conducted as previously described (Mayer *et al*. 2016). For OCTs and PMAT, their generation and maintenance followed similar procedures (Maier *et al*. 2021b). The cell lines were maintained in high glucose (4.5 g/L) and L-glutamine-containing DMEM, supplemented with 10% FBS, 1 µg/mL streptomycin, 100 IU/mL penicillin, and G418 (250 µg/mL) in a humidified atmosphere (37°C, 5% CO_2_) and in a subconfluent state. For 5-HT_2_ G protein dissociation assays, HEK 293T cells (ATCC) were used.

### 2.3. Uptake inhibition and release assays

Experiments were conducted in HEK293 cells as previously described (Mayer *et al*. 2016; Nadal-Gratacós *et al*. 2021), with minor modifications. Radiotracers were [^3^H]5-HT for SERT and [^3^H]MPP^+^ for DAT, OCT1-3, and PMAT. Non-specific uptake was determined in the presence of GBR12909 (50 μM) for DAT, paroxetine (3 μM) for SERT and decynium-22 (D22) (100 μM) for OCT1-3 and PMAT, and represented < 10% of total uptake.

### 2.4 Transporter electrophysiology: HEK293 cells and *Xenopus laevis* oocytes

HEK293 cells overly expressing the transporter of interest were used. For human DAT (hDAT), a stably expressing cell line was used (Giros *et al*. 1992; Sitte *et al*. 1998). For human SERT (hSERT), a GFP-tagged version of the transporter in a tetracycline-inducible construct was used as previously described (Hasenhuetl *et al*. 2018).

*Xenopus laevis* oocytes were injected with *in vitro* transcribed cRNA of either hDAT or hSERT. Subsequent electrophysiological studies were performed using the two-electrode voltage clamp technique (Oocyte Clamp OC-725; Warner Instruments, Hamden, CT, USA) (Vacca *et al*. 2022; Bhatt *et al*. 2022). The animal study was reviewed and approved by the Committee of the “Organismo Preposto al Benessere degli Animali” of the University of Insubria and nationally by Ministero della Salute (permit n. 449/2021-PR). The portions of ovary were used according to the Italian Law Art. 18 (3’R) 316 DLgs26_2014.

### 2.5. Gq dissociation bioluminescence resonance energy transfer (BRET): 5-HT_2A/2B/2C_ receptor activity

5-HT2 Gq dissociation BRET assays were performed as previously described (Cunningham *et al*. 2023; Lewis *et al*. 2023). HEK 293T cells (ATCC) were transfected in 10% dialyzed FBS (dFBS; Omega Scientific) in a 1:1:1:1 ratio of human receptor:Gαq-Rluc8:β3:GFP2-γ9 DNA constructs prepared in Opti-MEM (Invitrogen) using a 3:1 ratio of TransIT-2020 (Mirus Bio) µL:µg total DNA. Next day, cells were detached, centrifuged, resuspended and plated in 1% dFBS at an approximate density of 30 000 cells per well into poly-L-lysine-coated 96-well white assay plates (Greiner Bio-One). After approximately 24 hours, media was decanted and replaced with 60 µL per well of drug buffer (1× HBSS, 20 mM HEPES, pH 7.4), and incubated for at least 15 minutes at 37°C in a humidified incubator before receiving drug administration. Drug dilutions were made in drug buffer containing 0.3% BSA fatty acid free and 0.03% ascorbic acid. Drug dilutions were dispensed in 30 µL per well using multi-channels and plates were incubated at 37°C in a humidified incubator until reading. Next, plates were briefly taken out and coelenterazine 400a (5 µM final concentration; Nanolight Technology) was added 15 minutes before reading. After 60 minutes total time for incubation, plates were read in a PheraStarFSX or ClarioStar Plus (BMG Labtech; Cary, NC) at 1 second per well for at least 15 minutes for 3-5 cycles. BRET ratios of 510/400 luminescence were calculated per well and were plotted as a function of drug concentration. Data were normalized to % positive control (5-HT) stimulation and analyzed using nonlinear regression “log(agonist) vs. response” to yield E_MAX_ and EC_50_ parameter estimates.

### 2.6. Calcium flux activity of 5-HT_2A/2B/2C_ receptors by GCaMP6s fluorescence

To further study the interaction of our test compounds with 5-HT_2A_R and 5-HT_2B_R, HEK293 cells were generated expressing tetracycline inducible CFP-tagged versions of 5-HT_2A_ and 5HT_2B_ receptor and a constitutively expressing Ca^2+^ sensor GCaMP6s (Chen *et al*. 2013), as previously described (Mayer *et al*. 2023).

### 2.7. Computational pharmacology

#### 2.7.1 Protein and ligand structures preparation

In this study, we utilized the hSERT structure (PDB ID: 5I71; (Coleman *et al*. 2016)). To generate homology models of hDAT based on the hSERT structure, we employed MODELLER (Šali and Blundell 1993). Prior to molecular docking, both proteins were submitted to molecular dynamics simulations following the established protocol described in previous studies (Szöllősi and Stockner 2021; Gradisch *et al*. 2022).

#### 2.7.2. Molecular docking

The ligands were docked into the protein with the co-transported ions bound using the GOLD (Genetic Optimization for Ligand Docking) software version 2022.2.0 (Jones *et al*. 1997).

### 2.8. Hepatic metabolism

#### 2.8.1. Pooled human liver microsome/S9 fraction incubation for identification of phase I and II metabolites and isozyme mapping; LC-HRMS/MS conditions

Incubation using pooled human liver microsomes (pHLM) were prepared according to published procedures (Welter *et al*. 2013; Richter *et al*. 2016). ODMA, TDMA, or SeDMA were incubated with pooled human liver S9 fraction (pS9) (2 mg microsomal protein/mL) in accordance to a previous publication with minor modifications (Richter *et al*. 2017b). Incubation conditions for isozyme mapping followed an established protocol (Wagmann *et al*. 2016).

Regarding LC-HRMS/MS conditions, and according to previously published procedures, analyses were performed using a Thermo Fisher Scientific (TF, Dreieich, Germany) Dionex UltiMate 3000 RS pump consisting of a degasser, a quaternary pump, and an UltiMate Autosampler, coupled with a TF Q Exactive Plus equipped with a heated electrospray ionization (HESI)-II source (Gampfer *et al*. 2019).

### 2.9. Statistical analysis

Data plotting and statistical analyses were performed with GraphPad Prism 9 or 10 (GraphPad Software Inc., San Diego, CA, USA). For both batch and superfusion release assays, data were statistically analyzed using a mixed-effects model, employing Šidák’s correction for multiple comparisons. For the area under the curve (AUC) analyses, ordinary one-way ANOVA test for multiple comparisons was used. Both these two last statistical analyses explored possible significant differences between KHB and KHB+Mon (10 µM) conditions. Statistical significance was defined at a p-value lower than 0.05. *denotes p<0.05, **p<0.01 and ***p<0.001.

## 3. Results

### 3.1. MDMA and its analogs inhibit the [^3^H]substrate uptake at hSERT and hDAT at low micromolar concentrations in HEK293 cells

The interaction of MDMA with the monoamine transporters has been investigated extensively over the years (Rudnick and Wall 1992; Montgomery *et al*. 2007; Baumann *et al*. 2012; Sandtner *et al*. 2016; Shimshoni *et al*. 2017; Dolan *et al*. 2019; Luethi *et al*. 2019; Kolaczynska *et al*. 2022). Thus, to start our molecular characterization, we first explored the capability of MDMA and its analogs to inhibit the substrate uptake at hSERT and hDAT. The resulting uptake inhibition curves and the respective IC_50_ values were calculated (Fig. 1B; Table 1) and it was found that MDMA and its analogs interacted with hSERT and hDAT at low micromolar concentrations. Specifically, at hDAT, MDMA, ODMA, and TDMA had similar potencies inhibiting [^3^H]MPP^+^ uptake, resulting in virtually identical IC_50_ values, whereas SeDMA showed a slightly decreased inhibitory potency (IC_50_= 9.8 ± 2.2 µM), being about half as potent than its congeners. At hSERT, TDMA was 2-fold more potent (IC_50_= 6.4 ± 1.0 µM) than MDMA in inhibiting [^3^H]5-HT uptake, whereas ODMA was 2-fold less potent. In contrast, SeDMA displayed a similar inhibitory potency compared with MDMA. Due to each compound similar uptake inhibitory potencies at hSERT and hDAT, the calculated hDAT/hSERT ratios (Table 1) resulted in low values (<10), suggesting a low abuse liability for these compounds (Luethi and Liechti 2020).

### 3.2. MDMA and its analogs evoke robust [^3^H]5-HT release at hSERT, but moderate [^3^H]MPP^+^ release at hDAT

In order to establish the substrate vs. inhibitor profile of each experimental drug at hSERT and hDAT, the calculated IC_50_ values from the previous uptake inhibition assays were used in the subsequent release assays in HEK293 cells. In these experiments, the time-dependent efflux of [^3^H]5-HT through hSERT and of [^3^H]MPP^+^ through hDAT was evaluated in the presence or absence of monensin (10 µM). Monensin is an ionophore that dissipates the sodium gradient across cell membranes that selectively enhances the efflux caused by transporter substrates, which helps to distinguish them from non-transported inhibitors (Scholze *et al*. 2000). At hSERT, MDMA and the three test drugs elicited a significant release of [^3^H]5-HT in the presence of monensin (Fig. 1C), when compared with negative and positive controls (paroxetine (0.05 µM) and *para*-chloroamphetamine (*p*CA; 10 µM), respectively) (Fig. S1). The calculated area under the curve of the percentage of [^3^H]5-HT released between 8 and 18 minutes (AUC_8-18min_) revealed a statistically significant difference between the conditions without and with monensin for all the experimental compounds (e.g. (in µM) MDMA_KHB_ = 108.9 ± 34.0 vs. MDMA_MON_ = 282.3 ± 15.4) (Fig. 1E). On the other hand, at hDAT, MDMA and its analogs evoked a moderate release of [^3^H]MPP^+^ over time in the presence of monensin (Fig. 1D), compared with the negative and positive controls (GBR12909 (0.5 µM) and *(S)-*amphetamine (10 µM), respectively) (Supplementary Fig. 1C, 1D). This could be detected easier by the resulting AUC_8-18min_, which showed a significant difference between the conditions without and with monensin for any test drug (e.g. (in µM) MDMA_KHB_ = 33.3 ± 7.5 vs. MDMA_MON_ = 63.1 ± 9.3) (Fig. 1F). Altogether, these results suggest that all compounds act as substrates/ releasers at hSERT and hDAT, with more efficacy at hSERT.

### 3.3. hSERT- and hDAT-mediated currents confirm full and partial substrate profiles

To further differentiate the substrate profile of the experimental drugs, we performed electrophysiology using whole-cell patch-clamp configuration (V_h_=-60 mV) in HEK293 cells overexpressing either hSERT or hDAT. hSERT and hDAT are Na^+^ dependent transporters, and the application of a substrate (but not an inhibitor) elicits an inwardly-directed current that persists for the whole application. Thus, it is currently used to identify whether a test drug acts as a substrate (Bhat *et al*. 2017; Hasenhuetl *et al*. 2019). At hSERT, the application of MDMA analogs elicited dose-dependent inwardly directed currents that resembled those elicited by MDMA or 5-HT (Fig. 2A, B). Non-linear regression of the concentration-response curve led to EC_50_ values in the low micromolar range ((in µM) 5-HT: 0.17 < MDMA: 0.27 < TDMA: 0.44 < SeDMA: 0.77 < ODMA: 1.27), and similar E_MAX_ values (5-HT: 94%; MDMA: 92.6%; TDMA: 93.2%; SeDMA 97.1%; ODMA: 109.5%). At hDAT, instead, all the compounds elicited inwardly-directed currents (Fig. 2C, D), but their amplitude reached only 30-50% compared to the saturating concentration of DA (30 μM). Non-linear regression revealed EC_50_ values in the low micromolar range ((in µM) MDMA: 2.51 < ODMA: 3.97 < TDMA: 4.29 < SeDMA: 5.97 < DA: 6.40) and similar E_MAX_ values (MDMA: 31.9%; ODMA: 48.2%; TDMA: 30.1%; SeDMA: 25.7%), except for DA (E_max_: 104.1%). To rule-out that the effect was not due to system bias, we measured transporter-mediated currents in *Xenopus laevis* oocytes expressing hSERT or hDAT, as previously described (Sonders *et al*. 1997; Cao *et al*. 1998; Meinild *et al*. 2004). Fig. S2 shows the effects of either 5-HT/DA, MDMA, ODMA, and TDMA, on hSERT- and hDAT-mediated currents, respectively.

**Fig. 2.**
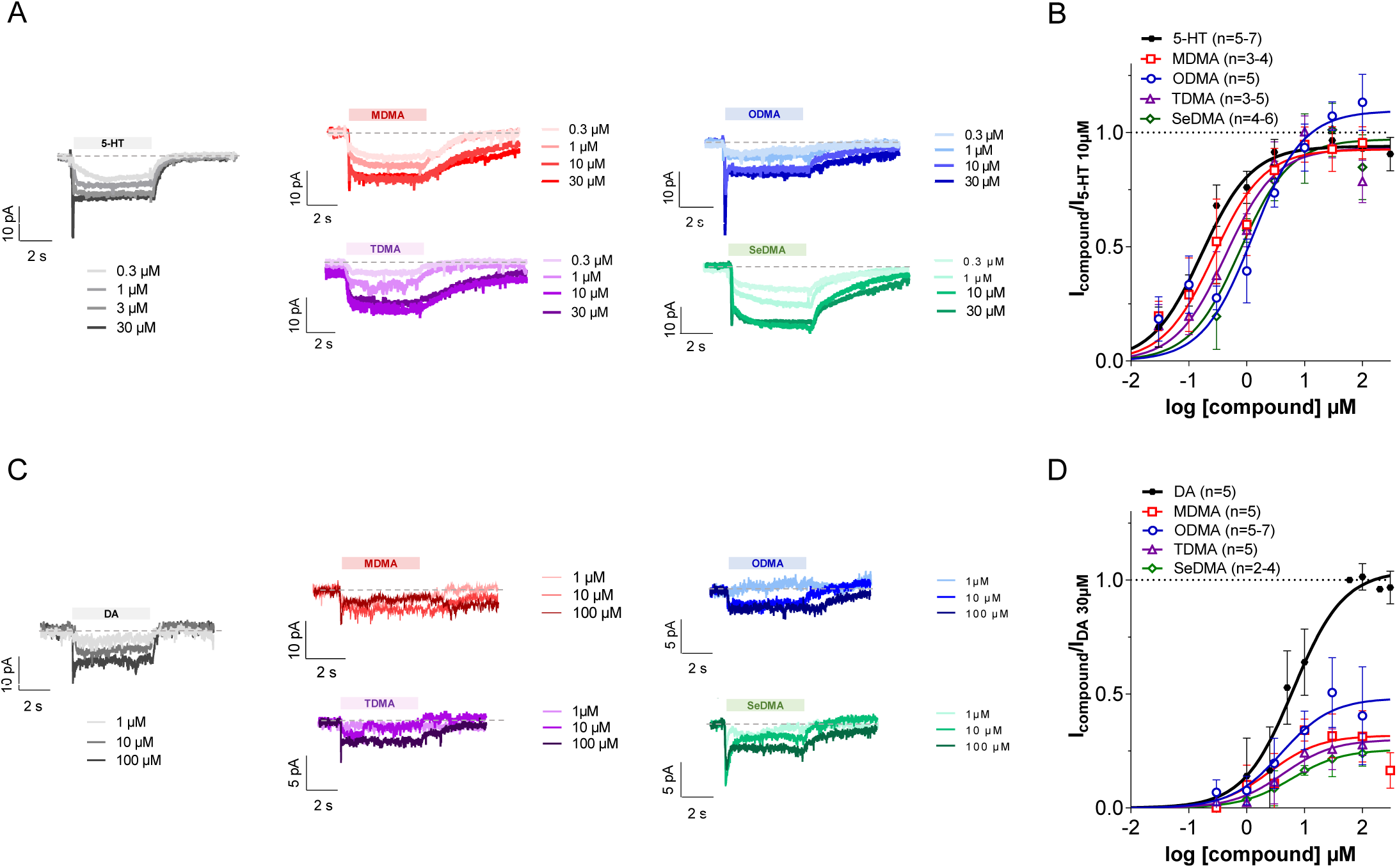
Measurement of transporter-mediated steady-state currents through whole-cell patch clamp (V_h_= −60 mV). (**A**) Representative single-cell traces showing hSERT-mediated currents elicited by increasing concentration of 5-HT (grey), MDMA (red), ODMA (blue), TDMA (purple), and SeDMA (green); (**B**) Concentration-response curves at hSERT. Data were normalized to the steady-state current of the saturating concentration of 5-HT (10 µM) and plotted using non-linear regression. The EC_50_ values were extrapolated (5-HT: 0.17 µM < MDMA: 0.27 µM < TDMA: 0.44 µM < SeDMA: 0.77 µM < ODMA: 1.27 µM) as well as the E_MAX_ values (5-HT: 94%; MDMA: 92.6%; TDMA: 93.2%; SeDMA 97.1%; ODMA: 109.5%); (**C**) Representative single-cell traces showing hDAT-mediated currents elicited by increasing concentration of DA (grey), MDMA (red), ODMA (blue), TDMA (purple), and SeDMA (green); (**D**) Concentration-response curves at hDAT. Data were normalized to the steady-state current of the saturating concentration of DA (30 µM) and plotted using non-linear regression. The EC_50_ values were extrapolated (MDMA: 2.51 µM < ODMA: 3.97 µM < TDMA: 4.29 µM < SeDMA: 5.97 µM < DA: 6.40 µM) as well as the E_MAX_ values (DA: 104.1%; MDMA: 31.9%; ODMA: 48.2%; TDMA: 30.1%; SeDMA: 25.7%). Data are mean ± SD from two to seven individual cells per each concentration.

Collectively, our results from these experiments revealed that MDMA and its analogs: 1) interacted with hSERT and hDAT at a similar low micromolar range; 2) elicited strong hSERT- and moderate hDAT-mediated efflux, and 3) accordingly showed full-efficacy for eliciting hSERT-mediated currents but partial-efficacy for eliciting hDAT-mediated currents. Taken together, the data support the conclusion that MDMA and its analogs show preference to act as full substrates at hSERT but as partial substrates at hDAT.

### 3.4. The binding poses of MDMA and its analogs overlap with the natural substrate, both at hSERT and hDAT

To better investigate the full vs. partial substrate dichotomy and the binding of MDMA, ODMA, TDMA, and SeDMA to hSERT and hDAT structures, we performed molecular docking calculations. Notably, these compounds exhibited remarkably similar binding poses when compared to each other (Fig. 3A, C), particularly when interacting with hSERT. The binding poses revealed that the positively charged amino group of the compounds faced the transmembrane helices forming the bundle domain (TM1, TM2, TM6, and TM7), while the aromatic ring system interacted with the scaffold domain (TM3, TM4, TM8, and TM9). This suggests that all ligands can interact with the same residues that interact with the endogenous substrates. Additionally, the observed conformations compare well with the 5-HT pose observed in hSERT structure (PDB ID: 7MGW; (Yang and Gouaux 2021)) and the poses of dopamine and methamphetamine observed in the drosophila dDAT structures (PDB ID: 4XP1 and 4XP6; (Wang *et al*. 2015)). Furthermore, upon analyzing the interacting residues (Fig. 3B, D (left and right panels); Fig. S3), we observed that these compounds established polar interactions through their charged amino group, as well as non-polar interactions through their aromatic ring system. The charged amine group formed electrostatic and hydrogen bond interactions with the side-chain of both D98 and S438, respectively, at hSERT and electrostatic and hydrogen bond interactions with the side-chain of D79 and the backbone of F320, respectively, at hDAT. Additionally, we noted that these compounds interacted with additional residues (marked with black stars) which have been shown to interact with 5-HT and dopamine (Yang and Gouaux 2021; Wang *et al*. 2015) in hSERT and hDAT, respectively (Fig. S3).

**Fig. 3.**
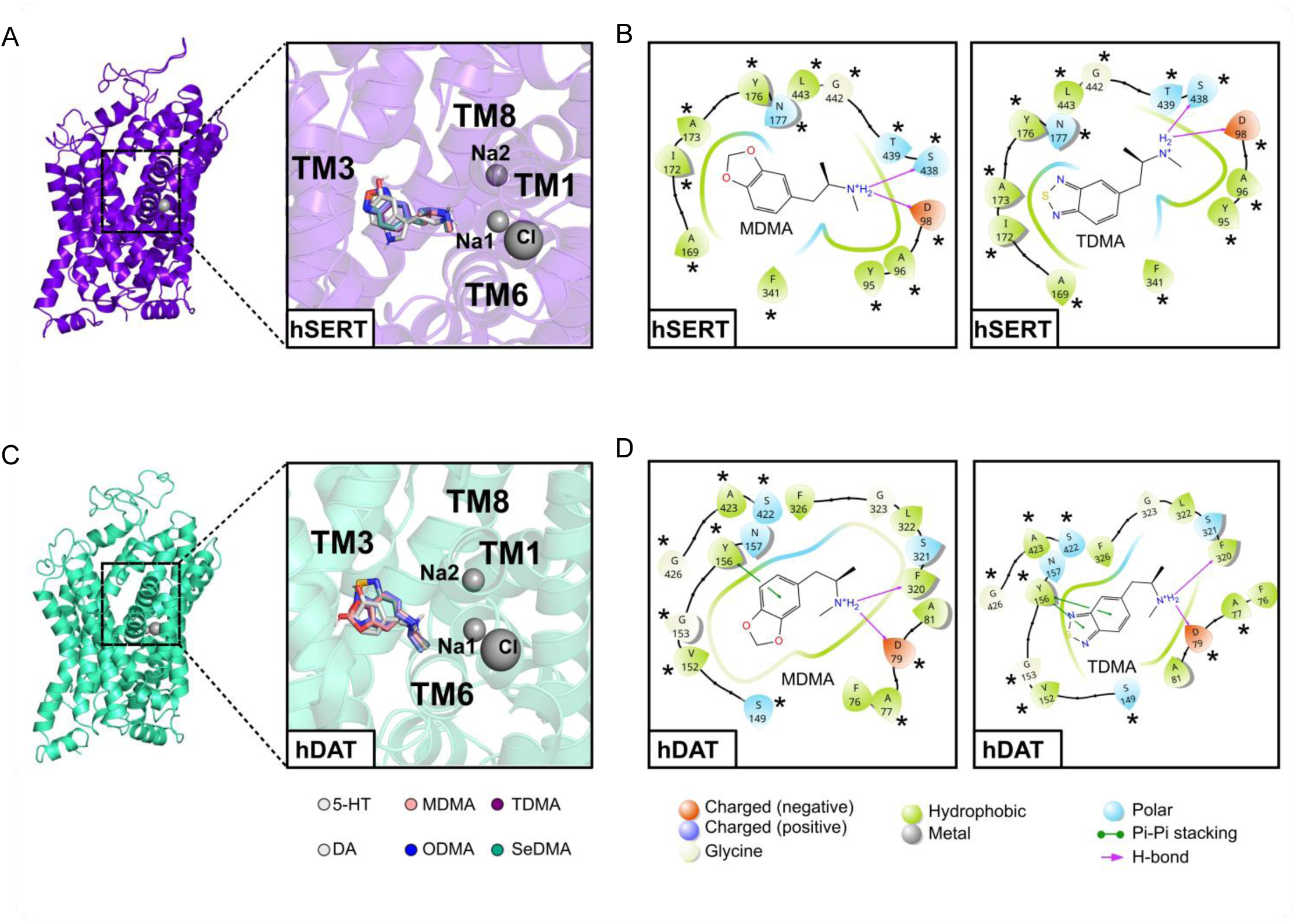
Molecular docking. (**A, C**) Outward-open structure of hSERT (purple) or hDAT (green cyan) and the binding poses of all ligands in each transporter. MDMA: salmon, OMDA: blue, TMDA: purple, SeDMA: green and both and serotonin (5-HT) and dopamine (DA) in light grey; (**B**) Left and right panels show the 2D interaction scheme of MDMA and TDMA molecules, respectively, highlighting the main interactions with hSERT (obtained using Maestro version 13.6.122); (**D**) Left and right panels show the 2D interaction scheme of MDMA and TDMA molecules, respectively, highlighting the main interactions with hDAT (obtained using Maestro version 13.6.122). The asterisks indicate the residues that also interact with 5-HT in hSERT (PDB ID:7MGW) and with DA in dDAT (PDB ID:4XP1). The residues of the protein are shown like guitar picks linked together on a string. The orientation of the guitar pick must be read as: the guitar picks pointing away from the ligand means the backbone of that residue is facing towards the ligand, and when the guitar pick is pointing toward the ligand that means the side chain of that residue facing the ligand.

### 3.5. MDMA activates 5-HT_2A_, 5-HT_2B_ and 5-HT_2C_ receptors more potently than its analogs

To further investigate our compounds directly at monoaminergic receptors, we explored their activity at 5-HT_2_ receptor subtypes, namely, 5-HT_2A_, 5-HT_2B_, and 5-HT_2C_ receptors. The activation of 5-HT_2A_R has been linked to the mechanism of action of psychedelics (McClure-Begley and Roth 2022). MDMA is known to be a weak 5-HT_2A_ receptor agonist (Nash *et al*. 1994), and this effect has been associated with its mesolimbic DA release and reinforcing properties (Teitler *et al*. 1990; Orejarena *et al*. 2011). Additionally, drugs causing valvular heart disease and primary pulmonary hypertension in humans have been found to share affinity for 5-HT_2B_ receptors (Rothman *et al*. 2000; Launay *et al*. 2002; Setola *et al*. 2003; Setola *et al*. 2005). MDMA has been shown to preferentially bind to and activate h5-HT_2B_ receptors with sub-micromolar affinity (Setola *et al*. 2003). Finally, 5-HT_2C_ agonists have been shown to decrease appetite (Thomsen *et al*. 2008).

To investigate the effect of MDMA and its analogs on 5-HT_2_ receptor activity, we measured Gq dissociation directly using a BRET-based assay system (Fig. 4A, B) (Cunningham *et al*. 2023; Lewis *et al*. 2023). Compared to MDMA, all three bioisosteres (ODMA, TDMA, and SeDMA) exhibited weaker potency and efficacy (Fig. 4C-F; Table 2) at 5-HT_2A/2B/2C_ Gq dissociation, with SeDMA being the weakest. In fact, all three bioisosteres were approximately 10-fold weaker to activate 5-HT_2B/2C_ receptors compared to MDMA. To complement these results, next we measured calcium flux responses using a GcAMP6 reporter which showed that all 3 bioisosteric MDMA analogs activate 5-HT_2A_ and 5-HT_2B_ receptors less efficaciously compared to MDMA (5-HT_2A_: E_MAX_=15.9%; 5-HT_2B_: E_MAX_= 15.4%) (Fig. 4S).

**Fig. 4.**
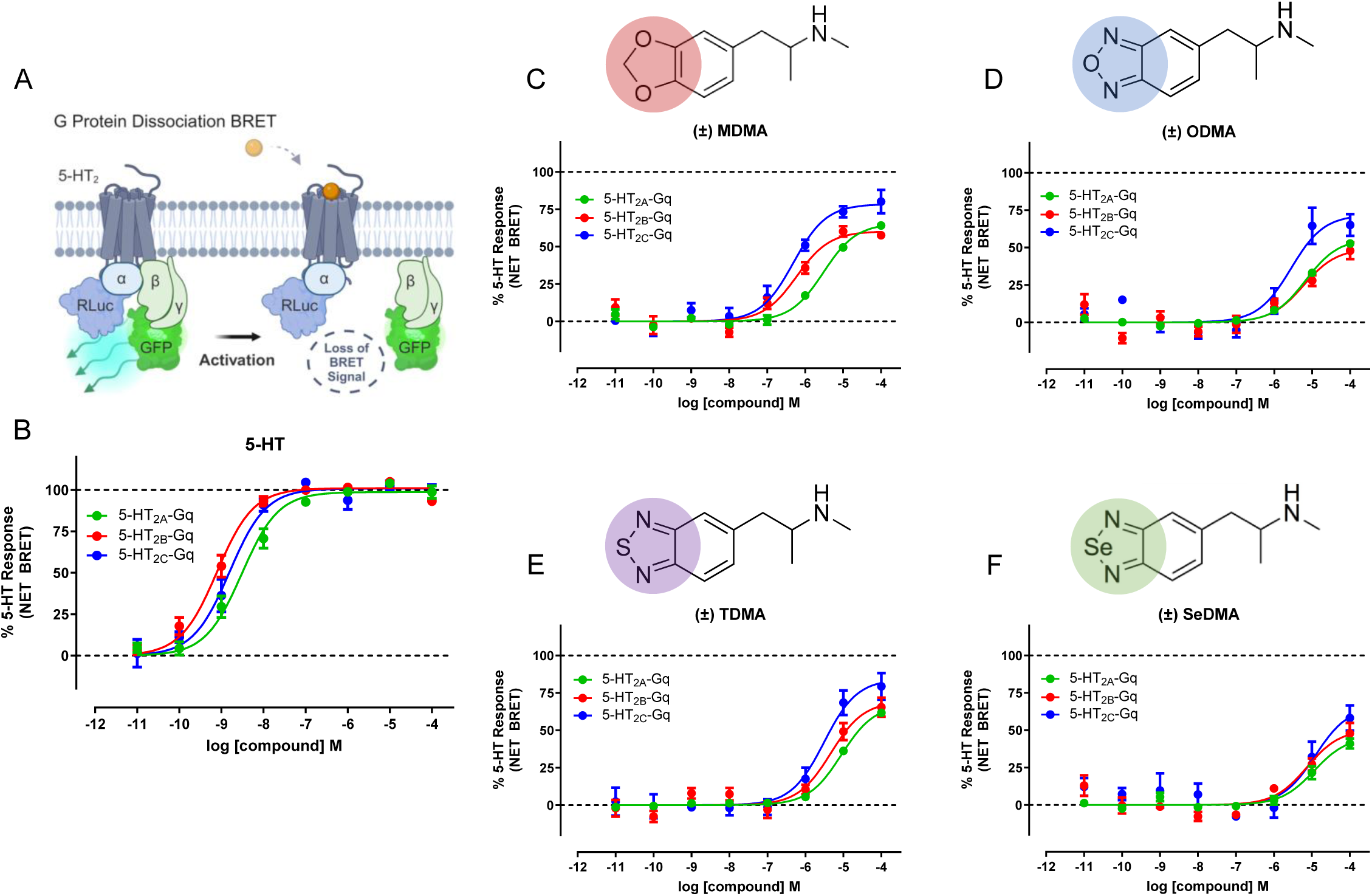
MDMA activates 5-HT_2A_, 5-HT_2B_, and 5-HT_2C_ receptors more potently than its analogs. (**A**) BRET Gq dissociation assay for human 5-HT2 receptors (5-HT_2A_-green, 5-HT_2B_-red, 5-HT_2C_-blue) using (B) 5-HT as positive control and measuring agonist activity of (C) MDMA compared to MDMA bioisosteric analogs (D) ODMA, (E) TDMA and (F) SeDMA. Data represent mean and SEM from three independent experiments.

**Table 2.**
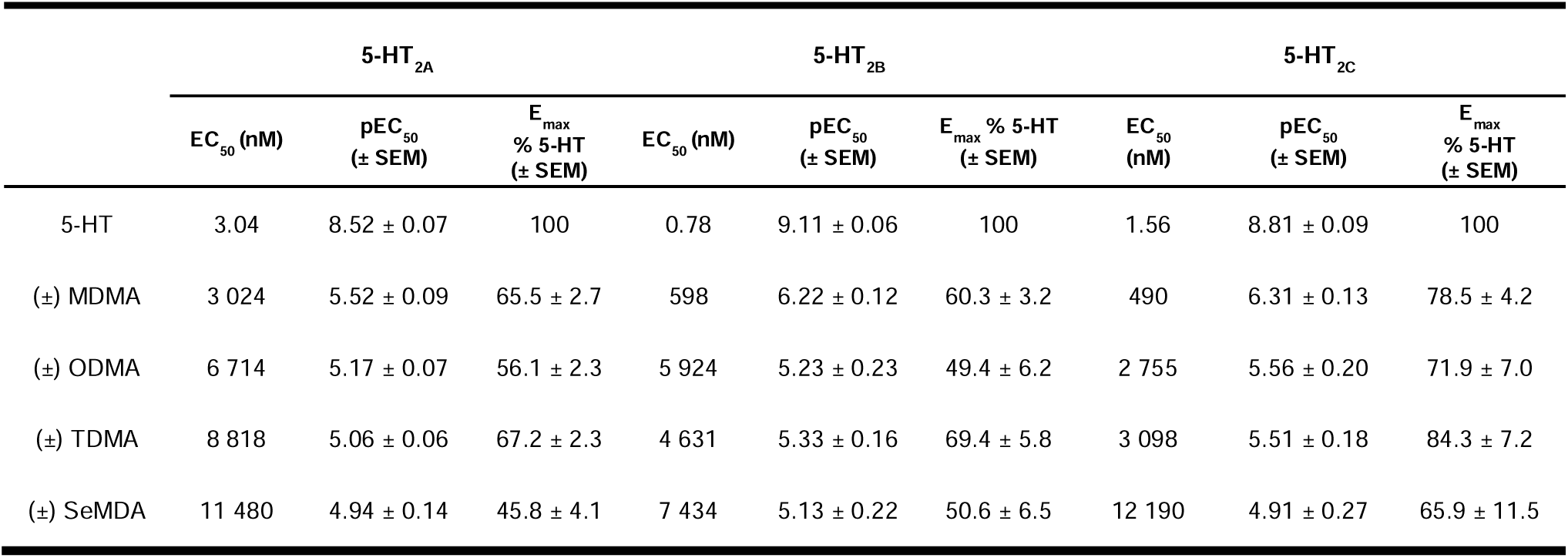
Gq dissociation BRET: 5-HT_2A/2B/2C_ receptor activity. 5-HT_2_ Gq dissociation EC_50_ and E_MAX_ parameter estimates of MDMA and analogs. 5-HT_2_ activation was measured using Gq/y9 dissociation by BRET. Data represent mean and SEM from three independent experiments performed in duplicate and reflect Fig. 4A-F.

### 3.6. CYP-mediated *N*-demethylation is the only hepatic metabolic pathway shared between MDMA and its analogs

After examining effects on the monoaminergic targets, the impact of the bioisosteric replacements involved in the design of the three MDMA analogs on *in vitro* hepatic metabolism was also investigated. Suitable *in vitro* systems can be used to mimic human metabolism. Such systems are pooled human liver microsomes (pHLM) combined with cytosol (pHLC) or pooled human liver S9 fraction (pS9), which are commonly used to identify phase I, but also phase II metabolites or both (Richter *et al*. 2017a). The S9 fraction usually contains cytosol and microsomes but the enzyme activities are usually lower than those of isolated microsomes or cytosol. Thus pHLM/pHLC is often tested besides pS9 (Richter *et al*. 2017a; Rock and Foti 2019; Brandon *et al*. 2003). All metabolites of ODMA, TDMA, and SeDMA detected from incubations with pHLM or pS9 mixtures along with their metabolite identification number, calculated exact mass of protonated molecule, elemental composition, and retention time are given in Table 3. Metabolites were tentatively identified by comparison of their HRMS^2^ spectra and fragmentation patterns of the parent compounds to those of the putative metabolites. All HRMS^2^ spectra of tentatively identified metabolites are shown in Fig. 5S (ODMA), 6S (TDMA), and 7S (SeDMA). For ODMA, three phase I metabolites were detected in all incubations. Metabolic reactions included *N*-dealkylation (ODMA-M1), *N*-hydroxylation (ODMA-M2), and hydroxylation (ODMA-M3). For TDMA, two phase I metabolites were found in all incubations. Hence, metabolic reactions included *N*-dealkylation (TDMA-M1) and *N*-hydroxylation (TDMA-M2). Finally, for SeDMA, one phase I reaction, namely an *N*-hydroxylation (SeDMA-M2) was found in all incubations. No phase II metabolites could be detected for any MDMA analogs. Thus, in this study, the main metabolic pathways of all three MDMA bioisosteres *in vitro* were *N*-demethylation and/or *N*-hydroxylation (Fig. 5A-5C). Comparing the results against the *in vitro* metabolism of MDMA (Richter *et al*. 2017a; Schwaninger *et al*. 2011a; Schwaninger *et al*. 2011b) (Fig. 5D), only *N*-demethylation was a common transformation. In contrast to MDMA, the bioisosteric replacement used in ODMA, TDMA, and SeDMA does not allow for a demethylenation to occur, which prevents catechol formation and subsequent formation of corresponding phase II metabolites as seen with MDMA.

**Fig. 5.**
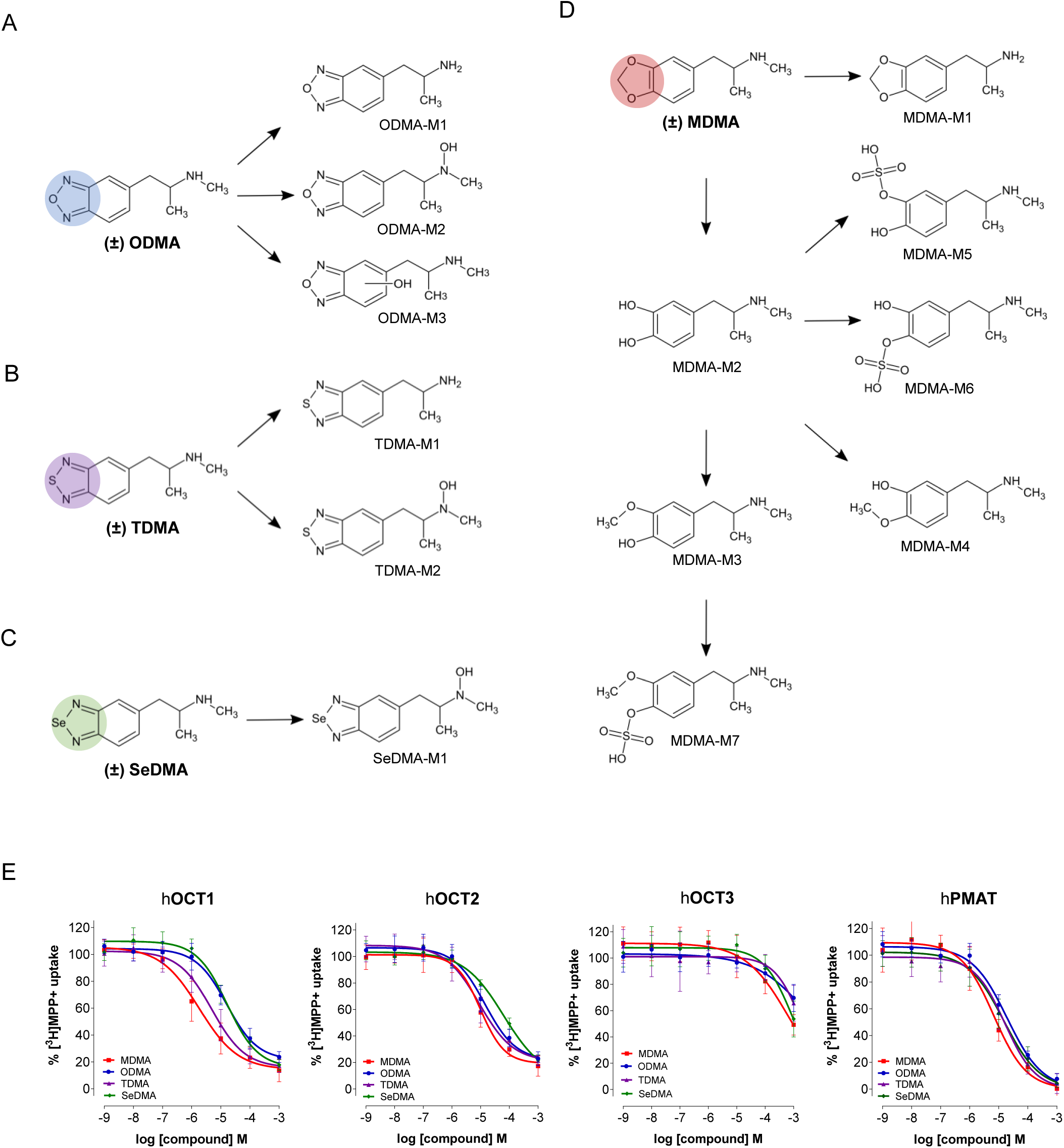
MDMA and its analogs differ in their hepatic metabolism and interact with the low-affinity high-capacity transporters OCT1-3 and PMAT at low micromolar concentrations. (**A-C**) Metabolic pathways of ODMA, TDMA, and SeDMA in incubations with pooled human liver microsomes. Undefined hydroxylation position is indicated by unspecific bonds. Metabolite-IDs correspond to Table 2; (**D**) Metabolic pathways of MDMA reported in literature in incubations with pooled human liver microsomes and/or S9 fraction (Richter *et al*. 2017a; Schwaninger *et al*. 2011b). Metabolite-IDs correspond to Table 2; (**E**) Uptake-inhibition curves at hOCT1, hOCT2, hOCT3, and hPMAT. Curves were plotted and fitted by non-linear regression, and data were best fitted to a sigmoidal dose-response curve to obtain IC_50_ values (see Table 1). Data are mean ± SD for individual experiments, performed in triplicate (n=3-4).

**Table 3.** Pooled human liver microsome/S9 fraction incubation for identification of phase I and II metabolites. Detection of ODMA, TDMA, and SeDMA and their phase I metabolites in pooled human liver microsomes and reported phase I and II metabolites of MDMA in literature in pooled human liver microsomes or S9 (Richter *et al*. 2017a; Schwaninger *et al*. 2011b) together with their metabolite identification numbers (ID), calculated exact mass of the protonated molecule (M+H+), elemental composition and retention time (RT). Metabolites were sorted by increasing mass. *, literature data, N.A., no retention time available for the used analytical method.

| Metabolite-ID | Metabolic reaction | Calculated exact mass, *m/z* | Elemental composition | RT, min |
| --- | --- | --- | --- | --- |
| ODMA | - | 192.1131 | C<sub>10</sub>H<sub>14</sub>N<sub>3</sub>O | 3.92 |
| ODMA-M1 | *N*-Dealkylation | 178.0975 | C<sub>9</sub>H<sub>12</sub>N<sub>3</sub>O | 4.56 |
| ODMA-M2 | *N*-Hydroxylation | 208.0935 | C<sub>10</sub>H<sub>14</sub>N<sub>3</sub>O<sub>2</sub> | 0.96 |
| ODMA-M3 | Hydroxylation at benzoxadiazole | 208.1081 | C<sub>10</sub>H<sub>14</sub>N<sub>3</sub>O<sub>2</sub> | 4.13 |
| TDMA | - | 208.0903 | C<sub>10</sub>H<sub>14</sub>N<sub>3</sub>S | 3.89 |
| TDMA-M1 | *N*-Dealkylation | 194.07464 | C<sub>9</sub>H<sub>12</sub>N<sub>3</sub>S | 4.16 |
| TDMA-M2 | *N*-Hydroxylation | 224.0852 | C<sub>10</sub>H<sub>14</sub>N<sub>3</sub>OS | 1.04 |
| SeDMA | - | 256.0347 | C<sub>10</sub>H<sub>13</sub>N<sub>3</sub>Se | 4.17 |
| SeDMA-M1 | *N*-Hydroxylation | 272.0297 | C<sub>10</sub>H<sub>14</sub>N<sub>3</sub>OSe | 1.16 |
| MDMA | - | 194.1176 | C<sub>11</sub>H<sub>15</sub>NO<sub>2</sub> | N.A. |
| MDMA-M1* | *N*-Dealkylation | 180.1019 | C<sub>10</sub>H<sub>13</sub>NO<sub>2</sub> | N.A. |
| MDMA-M2* | Demethylenation | 182.1176 | C<sub>10</sub>H<sub>15</sub>NO<sub>2</sub> | N.A. |
| MDMA-M3* | Demethylenation + methylation | 196.1332 | C<sub>11</sub>H<sub>17</sub>NO<sub>2</sub> | N.A. |
| MDMA-M4* | Demethylenation + methylation | 196.1332 | C<sub>11</sub>H<sub>17</sub>NO<sub>2</sub> | N.A. |
| MDMA-M5* | Demethylenation + sulfation | 262.0744 | C<sub>10</sub>H<sub>15</sub>NO<sub>5</sub>S | N.A. |
| MDMA-M6* | Demethylenation + sulfation | 262.0744 | C<sub>10</sub>H<sub>15</sub>NO<sub>5</sub>S | N.A. |
| MDMA-M7* | Demethylenation + methylation + sulfation | 274.0755 | C<sub>11</sub>H<sub>17</sub>NO<sub>5</sub>S | N.A. |

### 3.7. Different CYPs are involved in the transformation of MDMA analogs

The involvement of different cytochrome (CYP) 450 isozymes in the transformation of MDMA analogs was also analyzed. Mapping of isozymes is essential for predicting potential interactions, e.g., between drugs, or interindividual variations due to different expression of isozymes. Therefore, the involvement of 10 different CYP isozymes and FMO3 in the phase I biotransformation of ODMA, TDMA, and SeDMA was investigated using a monooxygenase activity screening. Results of isozyme mapping of initial phase I metabolites compared to pHLM incubations of ODMA, TDMA, and SeDMA are summarized in Table 4. The absence of interfering compounds was confirmed by blank incubations.

**Table 4.** Isozyme mapping of initial ODMA, TDMA, and SeDMA metabolites in comparison to pooled human liver microsomes (pHLM) and flavin-containing monooxygenase (FMO) 3 incubations. Metabolite-IDs correspond to Table 1. Isozyme mapping of MDMA metabolites reported in literature (Maurer *et al*. 2000; Meyer *et al*. 2008; Kraemer and Maurer 2002). Cytochrome P450 (CYP); *, literature data; +, detected; - not detected; /, not described in literature.

| Metabolite ID | pHLM | CYP |  |  |  |  |  |  |  |  |  | FMO |
| --- | --- | --- | --- | --- | --- | --- | --- | --- | --- | --- | --- | --- |
|  |  | 1A2 | 2A6 | 2B6 | 2C8 | 2C9 | 2C19 | 2D6 | 2E1 | 3A4 | 3A5 | 3 |
| ODMA |  |  |  |  |  |  |  |  |  |  |  |  |
| ODMA-M1 (N-Dealkyl) | + | + | - | + | - | - | + | + | - | + | - | - |
| ODMA-M2 (N-Hydroxy) | + | + | - | + | - | - | + | + | - | + | - | + |
| ODMA-M3 (Hydroxy) | + | - | - | - | - | - | - | + | - | + | - | - |
| TDMA |  |  |  |  |  |  |  |  |  |  |  |  |
| TDMA-M1 (N-Dealkyl) | + | + | - | + | - | - | + | + | - | + | - | - |
| TDMA-M2 (N-Hydroxy) | + | + | - | + | - | - | + | + | - | + | - | + |
| SeDMA |  |  |  |  |  |  |  |  |  |  |  |  |
| SeDMA-M1 (N-Hydroxy) | + | + | - | - | - | - | + | + | - | - | - | + |
| MDMA |  |  |  |  |  |  |  |  |  |  |  |  |
| N-Dealkyl* | + | + | - | + | - | - | + | + | - | + | + | / |
| Demethylenyl* | + | + | - | + | - | - | + | + | - | + | + | / |

### 3.8. MDMA analogs interact less potently with hOCT1, hOCT2, and hPMAT, compared with MDMA

In previous studies, we have found that a range of psychoactive substances differentially interact with the low-affinity high-capacity transporters (Mayer *et al*. 2018; Angenoorth *et al*. 2021; Maier *et al*. 2021a). Thus, to complement our studies on hepatic metabolism, we further investigated whether MDMA and its analogs were able to interact with the human organic cation transporters (hOCTs) 1 (hOCT1), 2 (hOCT2), and 3 (hOCT3), and human plasma membrane monoamine transporter (hPMAT). hOCT1-3, and hPMAT are generally involved in the uptake and elimination of various endogenous compounds, including monoamines, as well as of drugs, xenobiotics, and toxins. They are expressed both in peripheral organs (e.g. liver and kidney) and in the central nervous system, playing a major role in maintaining monoaminergic homeostasis (Koepsell 2020; Hayer-Zillgen *et al*. 2002). Thus, the interaction profile of the tested compounds was explored in hOCTs and hPMAT, and the respective IC_50_ values were calculated (Fig. 5; Table 1). In brief, MDMA and its analogs displayed IC_50_ values in the low micromolar range at hOCT1, hOCT2, and hPMAT, although SeDMA displayed a reduced inhibitory potential of the [^3^H]MPP^+^ uptake at hOCT-2 compared to its congeners (IC_50_ = 68.3 ± 30.5). Moreover, neither MDMA nor its congeners interacted with hOCT3 in a physiologically relevant concentration (IC_50_ values ≥ 696.2 ± 57.0 µM). Finally, at hPMAT, all compounds displayed IC_50_ values in the low micromolar range. In sum, MDMA analogs showed weaker interactions and displayed minor pharmacological differences in relation to uptake inhibition at hOCT1, hOCT2 and hPMAT, when compared with MDMA.

## 4. Discussion

Effective and long-lasting pharmacotherapies for specific neuropsychiatric disorders, including PTSD, continue to be important medical needs. Current therapies for PTSD include psychotherapy and/or pharmacotherapy, with selective serotonin reuptake inhibitors (SSRI) being the first-line agents (Williams *et al*. 2022). However, SSRIs show low efficacy in reducing PTSD symptoms severity (Hoskins *et al*. 2015; Williams *et al*. 2022). Recently, there has been a growing interest in psychedelics and other psychoactive drugs as possible therapeutic agents for the treatment of a range of psychiatric disorders (de Vos *et al*. 2021). In this context, MDMA is emerging as a candidate for the treatment PTSD in combination with psychotherapy (Mithoefer *et al*. 2018; Mithoefer *et al*. 2011; Mitchell *et al*. 2021; Mitchell *et al*. 2023).

In this study, we investigated three new MDMA bioisosteres and showed that they: 1) mimic MDMA interaction with the high-affinity low-capacity monoamine transporters hSERT and hDAT, displaying a full substrate profile at hSERT, but a partial substrate profile at hDAT; 2) are less potent and efficacious at 5-HT_2A/2B/2C_ receptors compared with MDMA; 3) differ from MDMA in their hepatic metabolism, sharing only *N*-demethylation during phase I without the formation of phase II metabolites; 4) interact slightly less potently with the low-affinity high-capacity transporters hOCT1, hOCT2, and hPMAT, compared with MDMA.

The therapeutic effects of MDMA involve a complex interplay between pharmacological and psychological effects. However, one of the key pharmacological mechanisms include the substrate-like activity and efflux-induction at monoamine transporters: MDMA is a substrate at SERT that leads to reverse the transport and potent 5-HT release both *in vitro* and *in vivo* (Rudnick and Wall 1992; Montgomery *et al*. 2007; Steinkellner *et al*. 2011; Sandtner *et al*. 2016). At DAT, most studies show that MDMA is also able to induce DA release both *in vitro* and *in vivo*, although this effect appears to be less pronounced (Baumann *et al*. 2012; Johnson *et al*. 1986). Regarding MDMA psychological effects, a part of its therapeutic action has been associated with its prosocial effects. Interestingly, in mice, the interaction of MDMA with SERT-containing 5-HT terminals in the nucleus accumbens has been demonstrated to be necessary and sufficient to explain this effect, whereas its non-social drug reward has been associated to the DA signaling in this same brain region (Walsh *et al*. 2018; Walsh *et al*. 2023; Heifets *et al*. 2019).

In the current study, we used heterologous expressing systems to evaluate the pharmacological mechanisms in greater detail and developed novel MDMA analogs. We have found that both MDMA and its analogs interacted with hSERT and hDAT at low micromolar concentrations with minor differences in their IC_50_ values (Table 1). Similar IC_50_ values have been reported for MDMA previously under comparable experimental conditions (Montgomery *et al*. 2007; Maier *et al*. 2018; Luethi *et al*. 2019; Ilic *et al*. 2020). Subsequent release assays showed that MDMA and its analogs induced significant SERT-mediated [^3^H]5-HT release and moderate hDAT-mediated [^3^H]MPP^+^ release. These data were further substantiated by whole-cell patch-clamp in HEK293 cells and two-electrode voltage-clamp in *Xenopus laevis* oocytes, which also confirmed that MDMA and its analogs elicit increasing concentration-dependent steady-state currents at hSERT and hDAT. Considering the amplitude of the currents at hSERT and hDAT, our findings suggest that MDMA and its analogs work as full substrates at hSERT, but as partial substrates at hDAT. Furthermore, molecular docking studies have shown that MDMA and its analogs display similar binding poses when binding at hSERT or hDAT (Fig. 3A, C). This result shows that the MDMA analogs preserve the same interaction pattern as MDMA despite the bioisosteric substitution of the methylenedioxy group. Interestingly, at hSERT, all interacting residues were shared between the compounds and 5-HT (Fig. 3B, Fig. S3). However, at hDAT, the compounds interacted with a larger number of residues compared to DA (Fig. 3D, Fig. S3, bottom panel). Such finding is particularly significant as it suggests that the docked compounds have a higher potential to mimic the binding of the native substrate in hSERT compared to hDAT, which is most likely a consequence of the higher structural similarity of these compounds to 5-HT as compared to dopamine, and supports our results with hSERT substrate preference over hDAT. The partial efficacy at hDAT might arise from slow binding kinetics, as shown already for the hSERT partial releaser PAL-1045 (Bhat *et al*. 2017). Further studies will be necessary to evaluate the extent to which binding kinetics might be involved in the pharmacodynamics of the novel MDMA analogs, and whether slow binding kinetics might play a role for their *in vivo* effects as shown already for a range of other psychoactive substances (Niello *et al*. 2023).

Given the unique psychopharmacological profile of MDMA, many other studies have explored the molecular pharmacology of other MDMA analogs, enantiomers, and/or metabolites at monoamine transporters (Montgomery *et al*. 2007; Sandtner *et al*. 2016; Shimshoni *et al*. 2017; Dolan *et al*. 2019; Pitts *et al*. 2018; Luethi *et al*. 2019; Kolaczynska *et al*. 2022). To our knowledge, this is the first study that investigated three novel MDMA analogs that reflected a bioisosteric replacement of the methylenedioxy group of MDMA and their impact on key molecular targets, together with a concomitant analysis of their hepatic metabolism.

In our study, these bioisosteric MDMA analogs exhibited less potent and efficacious agonist activity at 5-HT_2A_ and almost a 10-fold less potent activity at activating 5-HT_2B_ and 5-HT_2c_ receptors. For 5-HT_2A_, a weaker interaction with this receptor could translate in a reduced potential for mesolimbic DA release and reinforcing properties, together with a reduction/loss of the hallucinogenic potential of these compounds at higher doses (Teitler *et al*. 1990; Orejarena *et al*. 2011). Further studies will be necessary to confirm the reduced hallucinogenic effects *in vivo*, especially in relation to the 5-HT_2A_ signaling pathways that might be involved (Wallach *et al*. 2023). Importantly, the weaker agonism at 5-HT_2B_ receptor by the MDMA analogs suggests an improved pharmacological profile for the analogs with respect to MDMA, especially regarding the risk for cardiotoxicity. Nevertheless, the evaluation on risk for cardiotoxic effects with these novel MDMA bioisosteric analogs is warranted.

The hepatic metabolism studies showed that *N*-demethylation was the only metabolic pathway shared between MDMA and its analogs. The ring systems present in ODMA, TDMA, and SeDMA did not appear to undergo ring opening, in contrast to MDMA known to show demethylenation reactions and formation of catechol and subsequent phase II metabolites. Thus, our data support the hypothesis that the investigated MDMA analogs might be less likely to generate free radicals, in contrast with MDMA. Since demethylenation cannot occur, further studies are required to investigate whether this will impact on their pharmacokinetic properties. Regarding the isozyme mapping, several CYPs participated in the transformation of the MDMA analogs. Since ODMA and TDMA were mainly metabolized by CYP3A4, increased drug levels and intoxications may result from CYP3A4 inhibition, e.g., due to drug-drug or drug-food interactions (DDIs or DFIs, respectively). However, due to additional involvement of CYP2D6 in the metabolism of all three compounds, inhibition of CYP3A4 is expected to be less substantial in CYP2D6 poor metabolizers. In addition to CYP3A4 and CYP2D6, CYP1A2 was also involved in the phase I biotransformation of ODMA, TDMA, and SeDMA, which may prevent an increase in drug levels caused by CYP3A4 inhibitors or in CYP2D6 poor metabolizers. The *N*-demethylation catalyzed by CYP1A2 and CYP2D6 was also reported for MDMA (Maurer *et al*. 2000; Meyer *et al*. 2008; Kraemer and Maurer 2002). Since MDMA is known to be a substrate of CYP2D6 that can lead to its autoinhibition (De La Torre *et al*. 2004), further studies are encouraged to address this issue with these novel analogs to understand the potential of displaying DDIs or DFIs.

OCT1 and OCT2 play essential roles in drug pharmacokinetics, pharmacodynamics, and DDIs (Zhou *et al*. 2021), since they are involved in mediating hepatic uptake and renal secretion, respectively, of a wide range of cationic drugs (Suo *et al*. 2023). Overall, MDMA analogs demonstrated weaker interactions and minor pharmacological differences in relation to their uptake inhibition at OCT1, OCT2, and PMAT, compared with MDMA, which suggests potentially reduced risk for DDIs. None of the compounds interacted with OCT3 at relevant physiological concentrations.

Overall, these data suggest that ODMA, TDMA, and SeDMA are compounds that: 1) are capable of mimicking part of the MDMA molecular interactions at relevant physiological targets, such as hSERT and hDAT; 2) have lower activity at 5-HT_2A_,5-HT_2B_ and 5-HT_2C_ receptors, and 3) show improved metabolic properties. When compared with MDMA, further studies are needed to assess whether the favorable off-target profiles identified in this study translate to novel drug candidates with reduced side effect profiles.

## CRediT authorship contribution statement

**Ana Sofia Alberto-Silva:** Formal analysis, investigation, validation, visualization, writing – original draft, and writing – review & editing; **Selina Hemmer:** Formal analysis, investigation, validation, visualization, and writing – original draft; **Hailey A. Bock:** Formal analysis, investigation, validation, visualization, and writing – original draft; **Leticia Alves da Silva:** Formal analysis, investigation, validation, visualization, and writing – original draft; **Kenneth R. Scott:** Formal analysis, investigation, and methodology; **Nina Kastner**: Formal analysis and investigation; **Manan Bhatt:** Formal analysis and investigation; **Marco Niello:** Methodology, and writing - review & editing; **Kathrin Jäntsch:** Investigation; **Oliver Kudlacek:** Methodology, and writing - review & editing; **Elena Bossi**: Supervision, writing – review & editing, and funding acquisition; **Thomas Stockner:** Supervision, writing – review & editing, and funding acquisition; **Markus R. Meyer:** Supervision, and writing – review & editing; **John D. McCorvy:** Supervision, writing – review & editing, and funding acquisition; **Simon D. Brandt:** Conceptualization, and writing – review & editing; **Pierce Kavanagh:** Conceptualization, formal analysis, investigation, methodology, validation, and visualization; **Harald H. Sitte:** Conceptualization, Supervision, writing – review & editing, and funding acquisition.

## Data Availability

The data that support the findings of this study are available from the corresponding author upon reasonable request.

## Acknowledgments

We gratefully acknowledge the donation of the plasmids encoding for human DAT from the Caron Laboratory and human SERT from the Blakely Laboratory.

## Abbreviations

5-HT: 5-hydroxytryptamine; serotonin
BRET: Bioluminescence resonance energy transfer
CYP: Cytochrome P450
DA: Dopamine
DAT: Dopamine transporter
EC_50_: Half-maximal effective concentration
E_MAX_: Maximum effect
HEK293 cells: Human Embryonic Kidney 293 cells
IC_50_: Half-maximal inhibitory concentration
MDMA: 3,4-methylenedioxymethamphetamine
MPP^+^: 1-methyl-4-phenylpyridinium
NE: Norepinephrine
NET: Norepinephrine transporter
OCT: Organic cation transporter
*p*CA: *para*-chloroamphetamine
pHLC: Pooled human liver cytosol
pHLM: Pooled human liver microsomes
PMAT: Plasma membrane transporter
PTSD: Post-traumatic stress disorder
pS9: Pooled human liver S9 fraction
SERT: Serotonin transporter
SSRI: Selective serotonin reuptake inhibitor

