## Supplementary Information for "Bioisosteric analogs of MDMA with improved pharmacological profile"

Harald H. Sitte

Medical University Vienna

Center for Physiology and Pharmacology – Institute of Pharmacology

Währinger Straße 13a, 1090 Vienna Austria

T: +43-1-40160-31323

F: +43-1-40160-931300

### 1. Supplemental Materials and Methods

#### 1.1. Drugs and reagents

##### 1.1.1. Chemical synthesis of the experimental compounds ODMA, TDMA, and SedMA

*Generalized synthesis scheme*

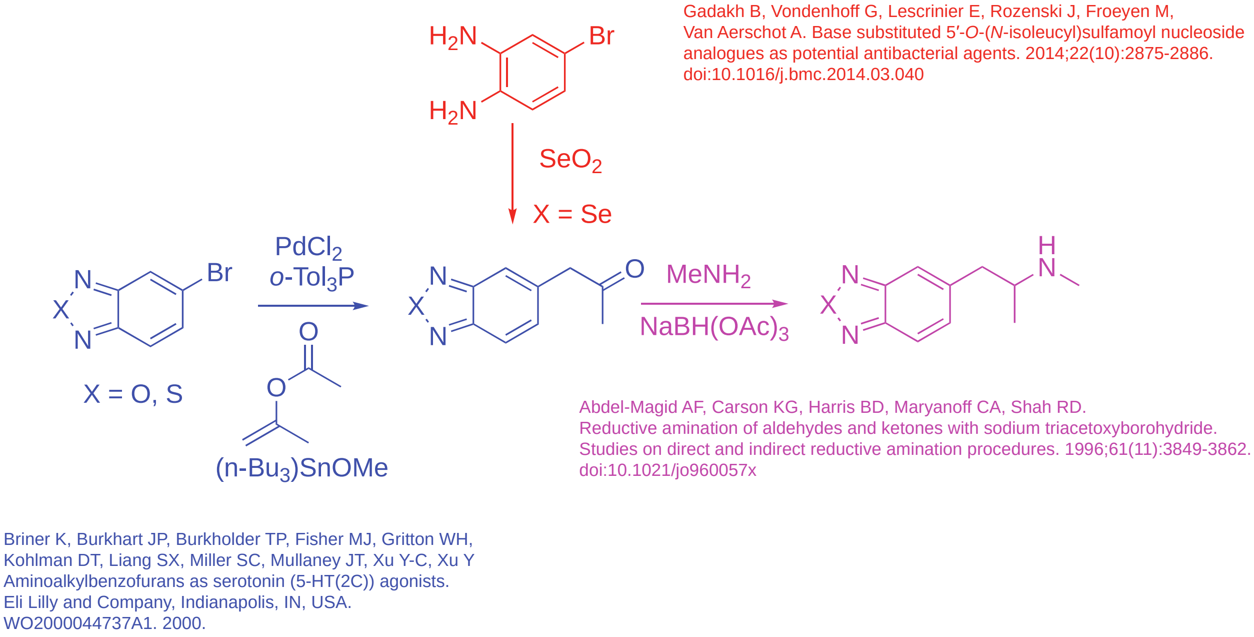

*1-(Benzo[c][1,2,5]oxadiazol-5-yl)-N-methylpropan-2-amine, hydrochloride salt (ODMA)*

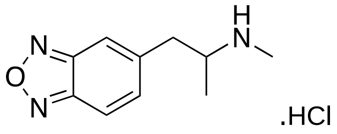

A mixture of 5-bromobenzo[*c*][1,2,5]oxadiazole (2.74 g, 13.8 mmol, Apollo Scientific), tri-(*o*-tolyl)phosphine (234 mg, 0.8 mmol) and tributyl tin methoxide (6.69 g, 6.0 mL, 20.8 mmol) in toluene (80 mL) was sparged with nitrogen for 15 min. *iso*-Propenyl acetate (2.8 g, 3 mL, 28 mmol) and palladium chloride (81 mg) were then added. The mixture was heated at 100°C for 5 h. After cooling to room temperature (RT), this was centrifuged (5 000 rpm/4330 *g* for 5 min) and the volatiles were removed from the supernatant to afford a brown oil. This was dissolved in dichloromethane (100 mL) and washed with aqueous sodium hydroxide (1 M, 4 x 100 mL). The organic layer was collected, dried (anhydrous magnesium sulphate) and evaporated to dryness to afford a dark heterogeneous oil. This was stored at 4°C overnight and then centrifuged (5000 rpm/4330 *g* for 5 minutes). The supernatant was removed and residue was washed with hexane to afford a dark reddish brown solid to give the corresponding ketone intermediate (1.473 g, 8.4 mmol, 61%).

This was added to ethanolic methylamine (8 M, 2.52 mL, 20.2 mmol) and glacial acetic acid (1.22 mL) in 1,2-dichloroethane (30 mL). The mixture was allowed to stir at RT for 1 h. Sodium triacetoxyborohydride (2.52 g, 11.9 mmol) was then added and stirring was continued at RT for 24 h. A mixture of water (50 mL) was added and the aqueous phase was made alkaline with and aqueous sodium hydroxide (50 % wt., 3 mL). The organic layer was collected and washed with water (50 mL) followed by brine (50 mL). Drying (anhydrous magnesium sulphate) followed by removal of the volatiles afforded a brown oil (1.296 g). A portion (421 mg) was purified by preparative thin layer chromatography (silica gel; 2 mm; ethyl acetate/methanol, 3/7), converted to the hydrochloride salt (2M hydrogen chloride in diethyl ether) and triturated with *tert*-butyl methyl ether to afford a light beige powder (134 mg + 360 mg = 504 mg). This was recrystallized from ethyl acetate/methanol to afford light beige powder (219 mg, 0.96 mmol, 7% from 5-bromobenzo[*c*][1,2,5]oxadiazole).

*1-(Benzo[c][1,2,5]thiadiazol-5-yl)-N-methylpropan-2-amine, hydrochloride salt (TDMA)*

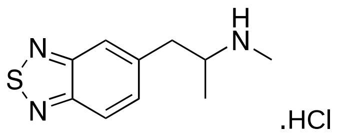

This was prepared in a similar manner to the procedure described above for ODMA starting with 5-bromobenzo[*c*][1,2,5]thiadiazole (3.22 g, 15 mmol). The intermediate ketone was obtained as a beige powder (1.076 g, 5.6 mmol, 37%). The crude freebase final product was obtained as reddish brown oil (1.189 g). The final product was obtained as a light beige powder (291 mg + 20 mg = 311 mg). This was recrystallized from ethyl acetate/methanol to afford colourless powder (207 mg, 0.85 mmol, 6% from 5-bromobenzo[*c*][1,2,5]thiadiazole).

*5-Bromobenzo[c][1,2,5]selenadiazole*

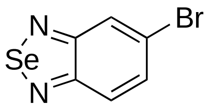

A mixture of 4-Bromo-1,2-diaminobenzene (6.0 g, 32 mmol) and selenium dioxide (4.0 g, 36 mmol) absolute ethanol (45 mL) was refluxed for 1 h. The solution was allowed to cool to RT and evaporated to dryness to afford a brown solid (7.7 g, 29 mmol, 91%).

*1-(Benzo[c][1,2,5]selenadiazol-5-yl)-N-methylpropan-2-amine, succinate salt (SeDMA)*

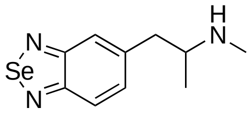

.succinate

This was prepared in a similar manner to the procedure described above for ODMA starting with 5-bromobenzo[c][1,2,5]selenadiazole (6.4 g, 24.24 mmol). The intermediate ketone was obtained as a brown powder (651 mg, 2.72 mmol, 11%). The freebase final product (212 mg, 0.83 mmol, 3% from 5-bromobenzo[c][1,2,5]selenadiazole) was dissolved in absolute ethanol (2 mL), succinic acid (50 mg, 0.42 mmol) was added and the salt was allowed to crystalize and then recrystallized from ethanol/ethyl acetate to afford an almost colourless powder (base:succinate mole ratio = 3:2).

Electron ionization mass spectra and nuclear magnetic resonance (NMR) spectra are shown at the end of the supplement.

##### 1.1.2. Other drugs and reagents

Acetonitrile (LC-MS grade), ammonium acetate, ammonium formate, formic acid (LC-MS grade), magnesium chloride ($MgCl_2$), methanol (LC-MS grade), and nicotinamide adenine dinucleotide phosphate ($NADP^+$) were obtained from VWR (Darmstadt, Germany). Acetyl coenzyme A (AcCoA), Acetyl-L-carnitine (ALC), carnitine acetyltransferase (CAT), dithiothreitol (DTT), reduced glutathione (GSH), isocitrate, isocitrate dehydrogenase, superoxide dismutase, 3'-phosphoadenosine-5'phosphosulfate (PAPS), potassium dihydrogenphosphate ($KH_2PO_4$), dipotassium hydrogenphosphate ($K_2HPO_4$), S-(5'-adenosyl)-L-methionine (SAM), superoxide dismutase (SOD), tris hydrochloride, and palmitic acid-$d_{31}$ were obtained from Sigma Aldrich (Taufkirchen, Germany). L-Tryptophan-$d_5$ and risperidone were purchased from Alsachim (Illkirch-Graffenstaden, France) and LGC Standards (Wesel, Germany).

The baculovirus-infected insect cell microsomes (Supersomes) containing 1 nmol/mL of human cDNA expressed cytochrome P450 (CYP) isoforms CYP1A2, CYP2A6, CYP2B6, CYP2C8, CYP2C9, CYP2C19, CYP2D6, CYP2E1 (2 nmol/mL), CYP3A4, CYP3A5 (2 nmol/mL), flavin-containing monooxygenase (FMO) 3 (5 mg/mL), pS9 (20 mg microsomal protein/mL), UGT reaction mixture solution A (25 mM UDP-glucuronic acid), UGT reaction mixture solution B (250 mM Tris HCl, 40 mM MgCl2, and 125 µg/mL alamethicin), pHLM (20 mg protein/mL) and pHLS9 (20 mg protein/mL) were obtained from Corning (Amsterdam, Netherlands). After delivery, the enzymes, pHLM, and pHLS9 were thawed at 37°C, aliquoted, snap-frozen in liquid nitrogen, and stored at -80°C until use.

#### 1.2. Uptake inhibition and release assays

In brief, HEK293 cells stably expressing the transporter of interest were seeded onto poly-D-lysine (PDL) coated 96-well plates (uptake-inhibition assays: 36 000 cells/well; batch release assays: 40 000 cells/well (hSERT) or 100 000 cells/channel (hDAT)) the day before the experiment. Uptake-inhibition assays: Prior to the uptake inhibition assay, DMEM was replaced with Krebs-HEPES buffer (KHB, composed of (in mM) 10 HEPES, 120 NaCl, 3 KCl, 2 $CaCl_2$, 2 $MgSO_4$, and 20 D-glucose, pH 7.3). The cells were pre-incubated with different concentrations of the drug in KHB at a final volume of 50 µL/well for 5 min (hSERT and hDAT) or 10 min (hOCT1-3, and hPMAT) at RT. Immediately after, the pre-incubation solution was removed and the cells were incubated with tritiated substrates: $[^3H]MPP^+$ (0.08 µM) for hDAT (1 min), $[^3H]$5-HT (0.1 µM) for hSERT (1 min), and [$^3$H]MPP$^+$ (0.05 µM) for hOCT1-3 and hPMAT (10 min), along with different concentrations of the drug in KHB. Finally, the [$^3$H]substrate was removed, and the cells were washed with ice-cold KHB. Afterwards, scintillation cocktail was added to each well and the multi-well plate was subjected to liquid scintillation counting to quantify tritium accumulated in each sample (Wallac 1450 MicroBeta® JET). Non-specific uptake was determined in the presence of paroxetine (3 µM) for hSERT, GBR12909 (50 µM) for hDAT, and decynium-22 (D22) 100 µM for hOCT1-3 and hPMAT and represented < 10% of total uptake. The uptake in the absence of the test compound was normalized to 100%, being the percentage of the uptake in the presence of different concentrations of the drug expressed as a percentage. Uptake-inhibition curves were plotted and fitted by non-linear regression, and data were best fitted to a sigmoidal dose-response curve to obtain $IC_{50}$ values. 1/hDAT $IC_{50}$:1/hSERT $IC_{50}$ formula was used to calculate hDAT/hSERT ratios. Batch release assays (hSERT): Prior to the release assay, DMEM was removed, and cells were preloaded with [$^3$H]5-HT (0.08 µM) (37°C, 5% $CO_2$, for 20 min). Subsequently, the cells were washed for 15 minutes four times using KHB at RT. Afterwards, in parallel and concomitant assays, cells were pre-incubated for 10 min in KHB or in KHB + Mon 10 µM. Finally, the substance of interest was added at a concentration close to its $IC_{50}$ previously determined in the uptake inhibition assays. The resulting supernatant was collected and transferred to a new well every 2 min. At least three independent experiments were performed in duplicate to each experimental drug. Liquid scintillation cocktail was added to the wells with remaining cells, to the wells with the transferred supernatant, and to the wells used for total uptake and total activity measurements (Wallac 1450 MicroBeta® JET). Total radioactivity present in the supernatant and in the remaining cells was set as 100%, and the amount of [$^3$H] substrate present in the supernatant was expressed as percentage of the total. Superfusion release assays (hDAT): A day prior to the experiment HEK293 cells stably expressing hDAT were seeded at a density of 100 000 cells/channel into 6 channel flow ibidi ® slides. Prior to the release assay, DMEM was removed, and cells were preloaded with [$^3$H]MPP$^+$ (0.05 µM) at 37°C, 5% $CO_2$, for 20 min. Subsequently, the slides were transferred to the superfusion apparatus, and the cells were washed for 15 minutes under constant flow of 0.5 mL/min using KHB at RT. Afterwards, a fraction was collected every two minutes in 8 mL vials containing 2 mL scintillation cocktail, starting with three KHB basal release fractions. This is followed by another 4 fractions of basal release with KHB with or without Mon (10 µM). Finally, the substance of interest was perfused at a concentration close to its previously determined $IC_{50}$ and five fractions were collected. Finally, cells were lysed using 1% SDS for the evaluation of the remaining radioactivity within the cells, and 3 fractions were collected. The measured amount of tritiated substrate in the respective fraction was expressed as percentage of tritiated substrate at the start of the fraction. To determine the specificity of drug-induced reverse transport, selective transporter inhibitors paroxetine (0.05 µM) (hSERT) and GBR12909 (0.5 µM) (hDAT) and effective releasers *p*CA (10 µM) (hSERT) and (S)-amphetamine (10 µM) (hDAT) were used.

**1.3. Transporter electrophysiology: HEK293 cells and *Xenopus laevis* oocytes**

HEK293 cells: 24 hours prior the experiment, cells were seeded at a low confluency in poly-D-lysine (PDL)-coated 3 cm Petri dishes, being kept at 37ºC and 5% $CO_2$ in a humidified atmosphere. On the day of the experiment, DMEM was replaced by extracellular solution composed of (in mM) 140 NaCl, 3 KCl, 2.5 $CaCl_2{\cdot}2H_2O$, 2 $MgCl_2{\cdot}6H_2O$, 20 D-(+)-glucose·$H_2O$ and 10 HEPES, pH = 7.4 adjusted with NaOH. Micropipettes (3.5–6 MΩ) were prepared from borosilicate glass capillaries (Science Products GmbH) using a P-97 puller (Sutter Instrument) and filled with an intracellular solution composed of (in mM) 133 K-gluconate, 5.9 NaCl, 1 $CaCl_2{\cdot}2H_2O$, 0.7 $MgCl_2{\cdot}6H_2O$, 10 HEPES, 10 EGTA, pH = 7.2, adjusted with KOH. Transporter-mediated currents were recorded in whole-cell voltage-clamp mode, at RT. This clamping was held at -60 mV using an Axopatch 700B amplifier and pClamp 11.2.2 software. Solutions were applied using a DAD-12 perfusion system connected to an 8-tube perfusion manifold (ALA Scientific Instruments). Current traces were filtered at 1 kHz and digitized at 10 kHz using a Digidata 1550 (MDS Analytical Technologies). Data analysis was performed using Clampfit 10.2 software (Molecular Devices, Sunnyvale, CA, USA). Transporter-mediated currents elicited by test drugs were normalized to the steady-state current amplitude elicited by a saturating concentration of the natural substrate 5-HT (10 µM) or DA (30 µM) applied to the same cell, to account for differences in cell expression.

*Xenopus laevis* oocytes: 1) oocytes preparation: *Xenopus laevis* adult females were anesthetized using tricaine methane sulphonate (MS222; 0.10% (w/v) solution in tap water) and by laparotomy the portions of the ovary were removed and treated with 1 mg/mL collagenase IA (Sigma Collagenase from Clostridium histolyticum) in calcium-free ND96 [96 mM NaCl, 2 mM KCl, 1 mM MgCl2, 5 mM HEPES; pH 7.6] for at least 1 h at 18 °C. Healthy and fully grown oocytes were manually selected in NDE solution (ND96 plus 2.5 mM pyruvate, 0.05 mg/mL gentamicin sulfate, and 1.8 mM CaCl2). After 24 hours, healthy-looking stage V and VI oocytes were micro-injected as previously described (Bhatt *et al.* 2022); 2) mRNA preparation: cDNAs were linearized (pNB1HsDAT and pOTVhSERT both with NotI), *in vitro* capped, and transcribed using 200 units of T7 RNA polymerase. All enzymes were supplied by Promega (Italy). *Xenopus laevis* frogs were maintained according to international guidelines (Delpire *et al.* 2011; McNamara *et al.* 2018). The animal study was reviewed and approved by the Committee of the "Organismo Preposto al Benessere degli Animali" of the University of Insubria and nationally by Ministero della Salute (permit nr. 449/2021-PR); 3) electrophysiology and data analysis: The controlling software was WinWCP version 4.4.6 (J. Dempster, University of Strathclyde, Glasgow, UK) or Clampex (Molecular Devices, Sunnyvale, USA). The borosilicate microelectrodes, with a tip resistance of 0.5–4 MΩ, were filled with 3 M KCl. Bath electrodes were connected to the experimental oocyte chamber via agar bridges (3% agar in 3 M KCl). The holding potential was kept at −60 mV for all the experiments. Transport-associated currents ($I_{tr}$) were calculated by subtracting the mean values in the absence of compound from those in its presence and normalized to the value of the natural substrate (5-HT (10 µM) for hSERT or DA (30 µM) for hDAT). The MDMA analogs have been tested at the concentration starting from 0.3 µM to 100 µM. Data analysis was performed using Clampfit 10.2 software (Molecular Devices, Sunnyvale, USA).

##### 1.4. Calcium flux activity of 5-HT$_{2A/2B/2C}$ receptors by GCaMP6s fluorescence

Cells were maintained in DMEM supplemented with 10% FBS, and, depending on expression vector used, G418 (250 µg/mL), zeocin (150 µg/mL), and/or blasticidin (6 µg/mL). 24h prior to experiments, cells were seeded in DMEM containing 2% FBS, with a density of 25 000 cells/well into poly-D-ornithine-coated black CulturPlate™ plates and tetracycline dependent expression was initiated by application of tetracycline (1 µg/mL). Prior to the experiments, DMEM was replaced by KHB for 1 hour until the beginning of the assay. The activation of the receptors was detected via changes in GCamP6S fluorescence as consequence of the increase of intracellular $Ca^{2+}$ concentration. Changes in fluorescence over time were measured with FlexStation 3 (Molecular Devices) plate reader (Ex 485 nm/ Em 525 nm). All experiments were conducted at RT. Data were normalized to the maximal fluorescence signal elicited by 5-HT (1 µM), 30 seconds after its application (maximal fluorescence value).

#### 1.5. Computational pharmacology

##### 1.5.1 Protein and ligand structures preparation

In brief, the proteins, along with the co-transporter ions (two $Na^+$ and one $Cl^-$) and the respective endogenous substrates (dopamine for hDAT and 5-HT for hSERT), were embedded in a membrane bilayer consisting of 70% 1-palmitoyl-2-oleoyl-sn-glycero-3-phosphocholine (POPC) and 30% cholesterol (CHOL) molecules. These systems were then equilibrated and simulated using Gromacs version 2019.3 (Abraham *et al.* 2015) and the amber99sb-ildn (Lindorff-Larsen *et al.* 2010) force field. After performing 100 nanoseconds of molecular dynamics simulations, we selected the equilibrated structures for each transporter to be used in subsequent docking calculations. The ligands were built as the (*R*)-enantiomer and with protonation for pH 7 by protonating the primary amino group of the ligands. With the exception of SeDMA, all of the ligands were built based on MDMA (obtained from the PUBCHEM databank (Kim *et al.* 2023)) using YASARA version 21.12.19 (Krieger and Vriend 2014). Next, we performed an energy minimization using YASARA to obtain the optimized geometries of the ligands. The calculations were carried out in vacuum using the AMBER 96 force field (Wang *et al.* 2004) with an 0.8 nm force cut-off and particle mesh Ewald algorithm to treat long range electrostatic interactions. After removing the structural stress by energy minimization, an annealing simulation was performed using a timestep of 2 fs with atom velocities scaled down by 0.9 every $10^{th}$ step until energy converged to less than 0.05 kJ/mol per atom during 200 steps.

The selenium atoms are not parameterized in the AMBER 96 force field but builded on the similarity in chemistry between sulphur and selenium. The sulphur containing TDMA structure was used to create the SeDMA by replacing the sulphur with selenium in the energy minimised TDMA.

##### 1.5.2. Molecular docking

For each system, 10 docking solutions have been generated. Calculations were performed using a cavity formed by a 0.6 nm radius sphere centered on the center of mass of the respective endogenous substrates (dopamine for hDAT and 5-HT for hSERT). The calculation was performed with a rigid receptor; therefore, the residues of the cavity were not allowed to rotate. The docking poses were ranked using the built-in GoldScore (Verdonk *et al.* 2003) scoring function. The number of Genetic Algorithm (GA) runs, and all other parameters were set as default. The selection of the GoldScore scoring function was based on re-docking calculations (Supplementary Figure 3A), where hSERT and hDAT were docked with their respective endogenous substrates, following the aforementioned protocol.

#### 1.6. Hepatic metabolism

##### 1.6.1 Pooled human liver microsome incubation for identification of phase I metabolites

According to published procedures (Welter *et al.* 2013; Richter *et al.* 2016), ODMA, TDMA, or SeDMA were dissolved freshly in methanol and subsequently diluted with 0.1 M phosphate buffer to obtain the required concentrations. Incubations were performed using a final concentration of 0 or 25 µM of the respective compound and 1 mg protein/mL pHLM at 37°C. The final incubation mixtures also contained 90 mM phosphate buffer, 5 mM isocitrate, 5 mM $Mg^{2+}$, 1.2 mM $NADP^{+}$, 200 U/mL superoxide dismutase, and 0.5 U/mL isocitrate dehydrogenase. A final incubation volume of 50 µL was obtained. The reaction was stopped after 60 min by adding 50 µL ice-cold acetonitrile containing L-tryptophan-$d_5$ (5 mg/L). Samples were centrifuged for 2 min at 18,407 *x g*. For each group, two replicates were prepared.

##### 1.6.2. Pooled human liver S9 fraction incubation for identification of phase I and II metabolites

According to a previous publication (Richter *et al.* 2017), 25 µg/mL alamethicin (UGT reaction mixture solution B), 90 mM phosphate buffer (pH 7.4), 2.5 mM $Mg^{2+}$, 2.5 mM isocitrate, 0.6mM $NADP^{+}$. 0.8 U/mL isocitrate dehydrogenase, 100 U/mL superoxide dismutase were preincubated for 10 minutes at 37°C. Thereafter, 2.5 mM UDP-glucuronic acid (UGT reaction mixture solution A), 40 µM PAPS, 1.2 mM SAM, 1 mM DTT, 10 mM GSH, and 2.5 µM substrate were added. The amount of organic solvent was below 1%

(Chauret *et al.* 1998). All given concretions are concentrations in final incubation mixture (final volume 300 µL). Reactions were started by adding ODMA, TDMA, or SeDMA. The maximum incubation time was 360 minutes and 30 µL aliquots were taken after 60 and 360 minutes. Reactions were terminated by adding 10 µL ice-cold acetonitrile containing tryptophan-d5 (5 mg/L) as internal standard (IS). Afterwards, tubes were cooled for 30 minutes at -20°C, centrifuged at 18,407 *x g* for 2 minutes, supernatants were transferred to autosampler vials and analyzed by LC-HRMS/MS. Blank incubation (without substrate) and control incubation (without S9) were done to confirm the absence of interfering compounds and to identify not metabolically formed compounds. All incubations were performed in duplicate.

**1.6.3. Isozyme mapping**

According to an established protocol (Wagmann *et al.* 2016), ODMA, TDMA, or SeDMA were incubated at a final concentration of 25 µM with 50 pmol/ml of CYP1A2, CYP2A6, CYP2B6, CYP2C8, CYP2C19, CYP2D6, CYP2E1, CYP3A4, and CYP3A5, respectively, 0.25 mg protein/mL FMO3, or 1 mg microsomal protein/mL pHLM. Furthermore, incubation mixtures contained following components: isocitrate (5 mM), isocitrate dehydrogenase (0.5 U/mL), $MgCl_2$ (5 mM), $NADP^+$ (1.2 mM), 90 mM phosphate buffer (pH 7.4), and superoxide dismutase (200 U/mL). According to the manufacturer recommendation, incubations with CYP2A6 and CYP2C9 were conducted by replacing phosphate buffer with Tris buffer. Incubations (50 µL final volume) were performed at 37°C for 30 min and terminated by adding 50 µL of ice-cold acetonitrile. Afterwards, mixture was centrifuged at 18 407 *x g* for 5 min, supernatants were transferred into autosampler vials, and 1 µL was injected onto LC-HRMS/MS system. Incubations with pHLM were used as positive controls. Negative controls (without enzyme) were prepared to identify not metabolically formed compounds. All incubations were done in duplicate.

**1.6.4. LC-HRMS/MS conditions**

Hydrophilic interaction chromatography (HILIC) elution was performed using a Merck (Darmstadt, Germany) SeQuant ZIC HILIC (150 mm x 2.1 mm, 3.0 µm). The mobile phase consisted of aqueous ammonium acetate (200 mM, eluent A) and acetonitrile containing formic acid (0.1%, *v/v*, eluent B). The flow rate was set to 500 µL/min using the following gradient: 0-1min hold 2% A, 1-5 min to 20% C, 5-8.5

min to 60% A, 8.5-10 min hold 60% A, and 10-12 min hold 2% A. Injection volume was set to 5 µL for all samples. For preparation and cleaning of the injection system, isopropanol:water (90:10, *v/v*) was used. The following settings were used: wash volume, 100 µL; wash speed, 4000 nL/s; loop wash factor, 2. Column temperature for every analysis was set to 40°C. HESI-II source conditions were as follows: ionization mode, positive; sheath gas, 60 AU; auxiliary gas, 10 AU; sweep gas, 3 AU; spray voltage, 3.5 kV; heater temperature 320°C; ion transfer capillary temperature, 320°C; and S-lens RF level, 50.0. Mass spectrometry was performed using full scan and subsequently data-dependent acquisition (DDA) with priority to mass-to-charge ratios (*m/z*) of parent compounds and their expected metabolites. The settings for full scan data acquisition were as follows: resolution 35,000 at *m/z* 200; microscan, 1; automatic gain control (AGC) target, 1e6; maximum injection time, 120 ms; scan range, *m/z* 50-750; spectrum data type; centroid. Settings for DDA mode with an inclusion list containing the monoisotopic masses of ODMA, TDMA, or SeDMA and their expected metabolites were as follows: resolution, 17 500; microscans,1; isolation window, 1.0 *m/z*; loop count, 5; AGC target, 2e5; maximum IT, 250 ms; dynamic exclusion, 5 seconds; option "pick others" enabled; high collision dissociation cell with stepped normalized collision energy, 17.5, 35.0, 52.5; exclude isotopes, on; spectrum data type, profile; and underfill ratio, 1%. Inclusion list contained *m/z* values of likely formed metabolites such as hydroxy or *N*-dealkyl metabolites (phase I) as well as sulfates or glucuronides (phase II), and combinations of them. ChemSketch 2012 12.01 (ACD/Labs, Toronto Canada) was used to draw structures of hypothetical metabolites and to calculate exact masses. TF Xcalibur software version 4.5.474.0 was used for data handling.

#### 2. Supplementary figures

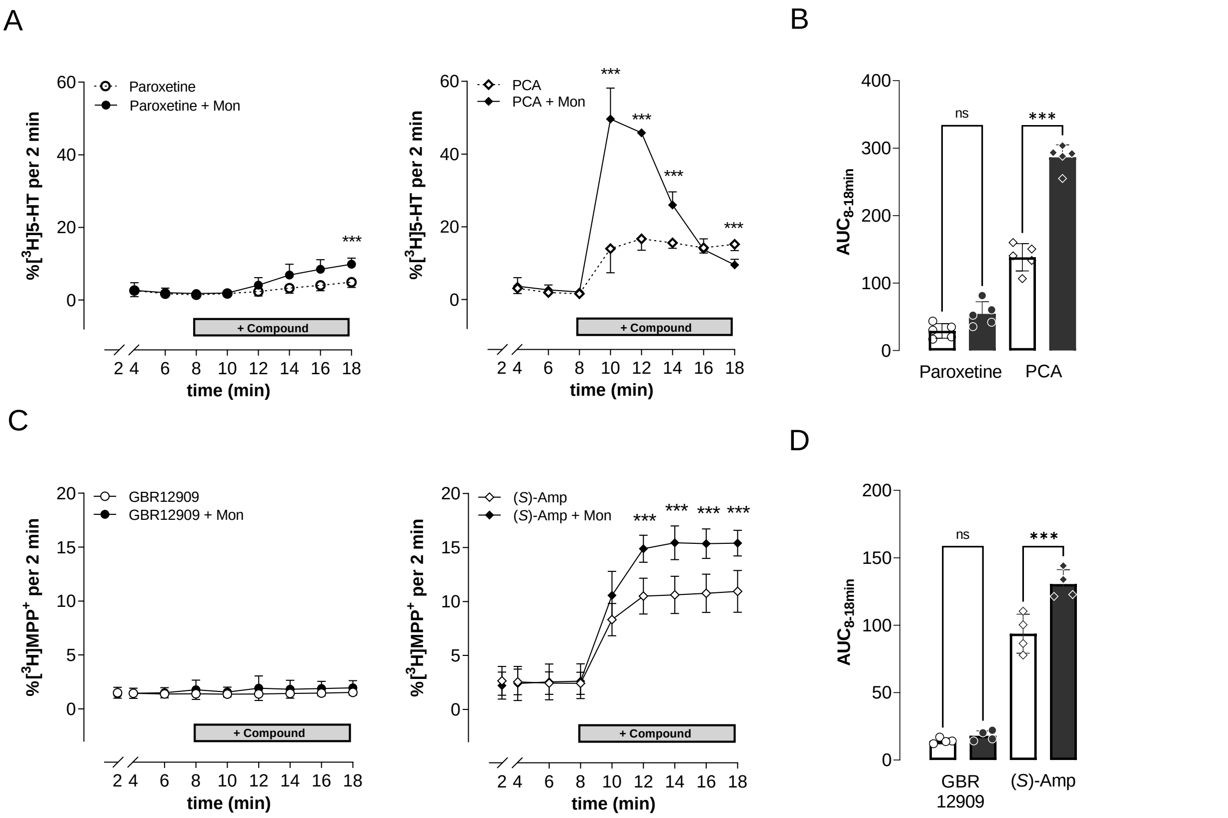

**Figure S1. hSERT- and hDAT-mediated efflux when exposed to a known transporter inhibitor or substrate/releaser.** (**A**) Effects of paroxetine and *p*CA on transporter-mediated release (batch) of preloaded [$^3$H]5-HT from HEK293 cells stably expressing hSERT. As indicated, after 8 min of basal release (either in KHB or KHB+Mon (10 µM)), the negative or positive control compounds were added (paroxetine (0.05 µM) and *p*CA (10 µM), respectively). (**B**) Calculated $AUC_{8\text{-}18min}$ of the [$^3$H]5-HT released by each compound in KHB (-) or KHB+Mon (+) conditions ($paroxetine_{KHB}$ = 29.2 ± 10.9 vs. $paroxetine_{MON}$ = 54.3 ± 18.0; $pCA_{KHB}$ = 138.2 ± 20.3 vs. $pCA_{MON}$ = 286.4 ± 18.5). (**C**) Effects of GBR12909 and *(S)*-amphetamine on transporter-mediated release (superfusion) of preloaded [$^3$H]$MPP^+$ from HEK293 cells stably expressing hDAT. As indicated, after 8 min of basal release (either in KHB or KHB+Mon (10 µM)), the negative or positive control compounds were added (GBR12909 (0.5 µM) and *(S)*-amphetamine (10 µM), respectively). (**D**) Calculated $AUC_{8\text{-}18min}$ of the [$^3$H]$MPP^+$ released by each compound in KHB (-) or KHB+Mon (+) conditions ($GBR12909_{KHB}$ = 14.2 ± 2.0 vs. $GBR12909_{MON}$ = 18.0 ± 3.7; *(S)*-$amphetamine_{KHB}$ = 93.7 ± 14.5

vs. *(S)*-amphetamine$_{MON}$ = 130.5 ± 10.7); Data are mean ± SD for individual experiments, performed in duplicate (n=5) (batch release assays) or in triplicate (n=4) (superfusion release assays). Statistical analyses explored possible significant differences between KHB and KHB+Mon conditions. Statistical significance was defined at a p-value lower than 0.05. *denotes $p<0.05$, **$p<0.01$ and ***$p<0.001$.

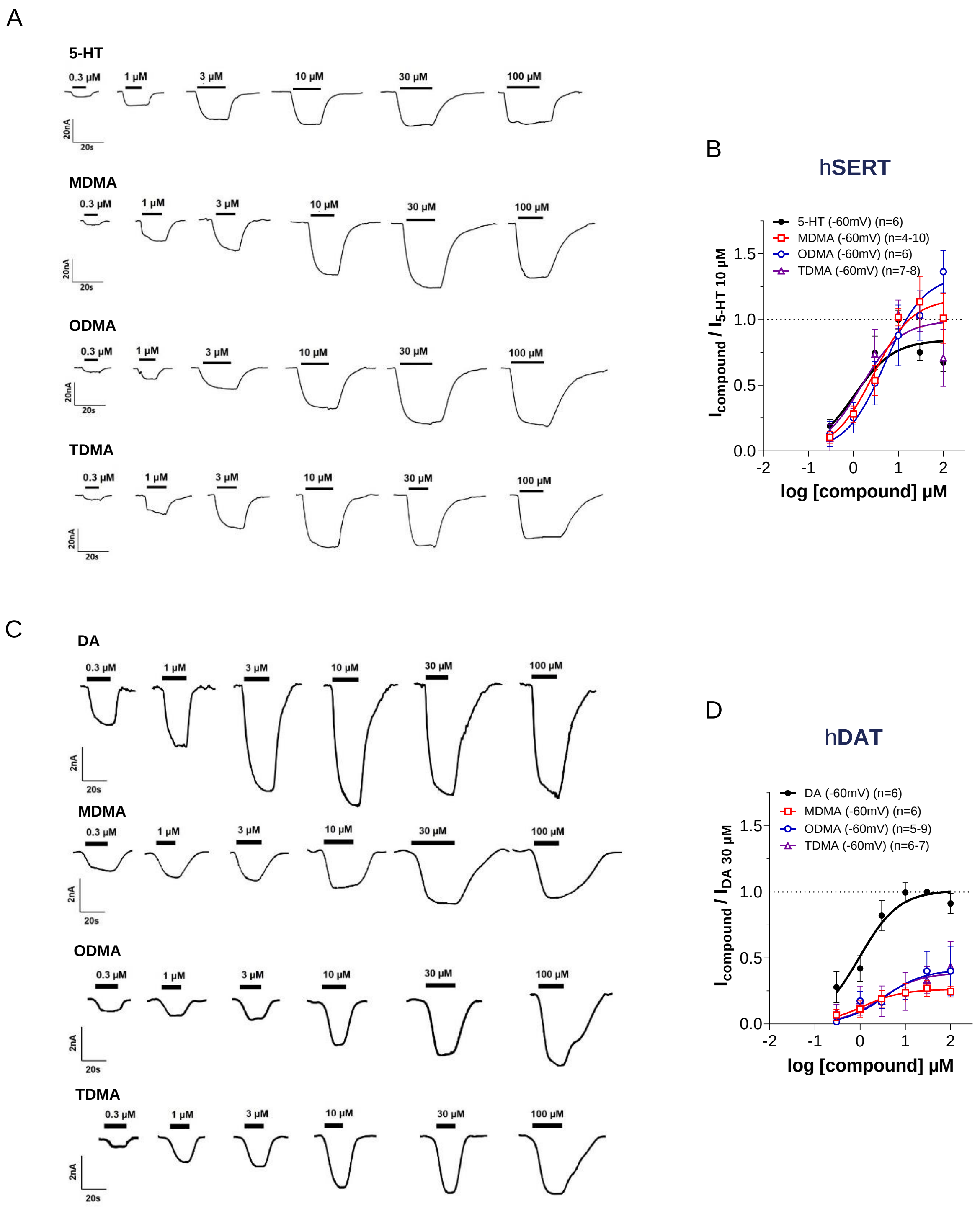

**Figure S2. Measurement of transporter-mediated inward currents through two-electrode voltage-clamp ($V_h$ = -60 mV).** (**A**) Representative single-oocyte traces showing hSERT-mediated currents elicited by increasing concentration of 5-HT (upper panel) and MDMA (bottom panel); (**B**) At hSERT, 5-HT, MDMA, ODMA, and TDMA evoked robust concentration-related inward currents that followed a bell-shaped concentration response. The greatest magnitude of current produced by MDMA and its analogs was equivalent to that produced by 5-HT (10 µM). Thus, 5-HT, MDMA, and its analogs displayed similar $EC_{50}$ values (5-HT: 1.05 µM < TDMA: 1.51 µM < MDMA: 2.77 µM < ODMA: 5.08 µM); (**C**) Representative single-oocyte traces showing hDAT-mediated currents elicited by increasing concentration of 5-HT (upper panel) and MDMA (bottom panel); (**D**) At hDAT, MDMA and its congeners could only evoke 30-40% of the inward currents elicited by DA (30 µM), nevertheless showing similar $EC_{50}$ values (DA: 0.97 µM < MDMA: 1.13 µM < TDMA: 2.90 µM < ODMA: 3.47 µM). Data were normalized to the steady-state amplitude current of 5-HT (10 µM) or DA (30 µM) and plotted using non-linear regression. Data are mean ± SD (n represents the number of individual oocytes which were used for each compound and concentrations tested).

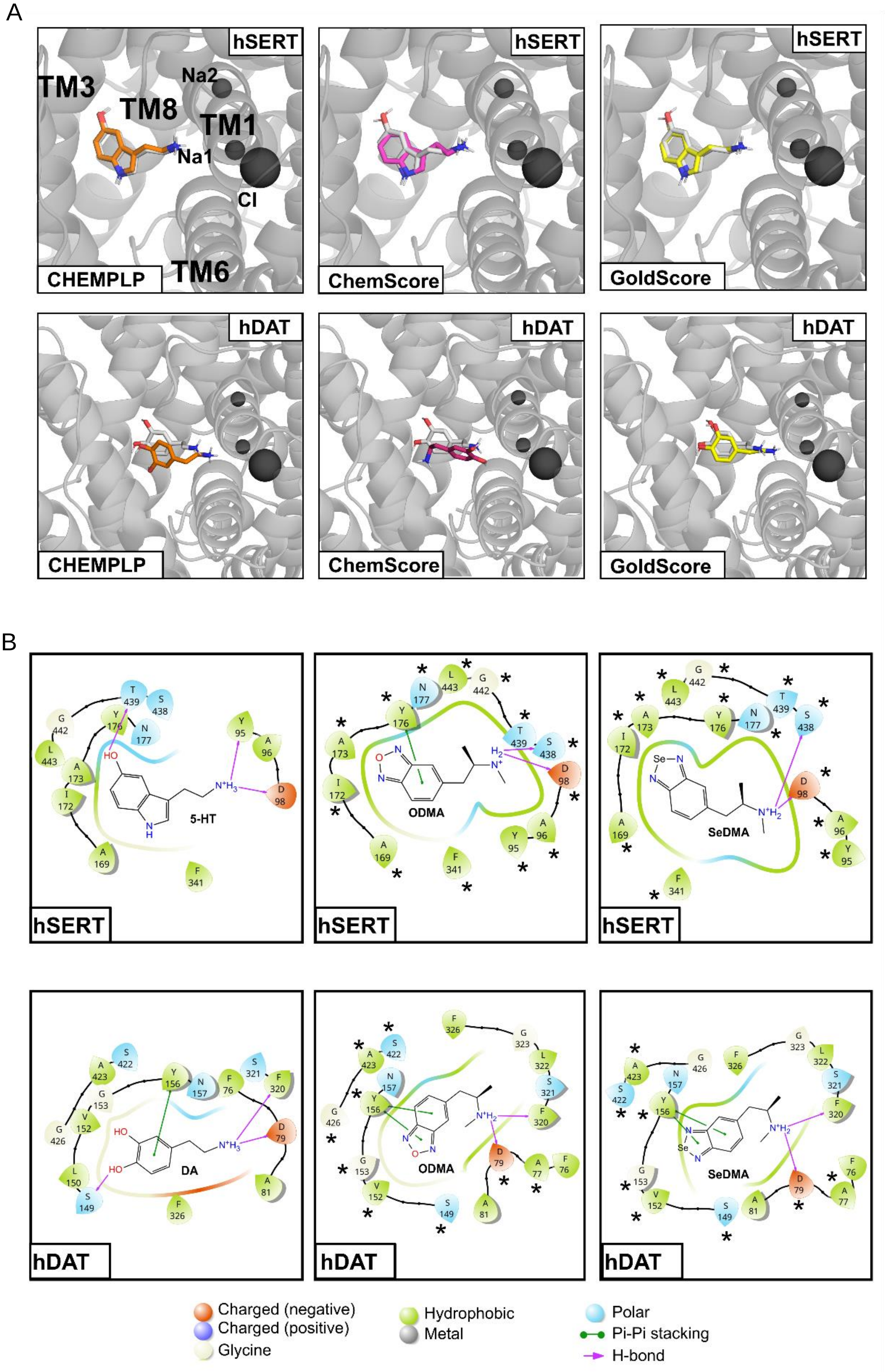
A
hSERT
TM3
Na2
TM8
TM1
Na1
Cl
TM6
CHEMPLP
ChemScore
GoldScore
hDAT
B
5-HT
ODMA
SeDMA
DA
Charged (negative)
Charged (positive)
Glycine
Hydrophobic
Metal
Polar
Pi-Pi stacking
H-bond

**Figure S3. Molecular docking**. (**A**) Re-docking of 5-HT (upper panel) in hSERT and DA in hDAT (lower panel) and their respective top-1 binding poses ranked based on three different scoring function (GoldSore, CHEMPLP and ChemScore). Both 5-HT and DA are colored in white/grey and the predicted binding poses are colored in orange, magenta or yellow; (**B**) 2D interaction scheme, obtained using Maestro version 13.6.122, of 5-HT or DA (left panels), ODMA (middle panels), and SeDMA (right panels) molecules, respectively, highlighting the main interactions with both hSERT (upper panels) and hDAT (lower panels). The asterisks indicate the residues that also interact with dopamine in dDAT (PDB ID:4XP1) and serotonin in hSERT (PDB ID:7MGW).

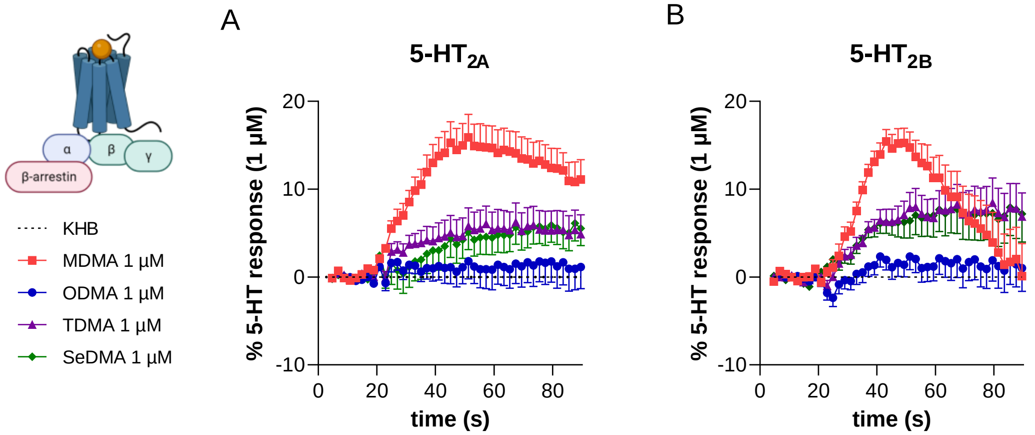

**Figure S4**. **Effects of 1 µM of MDMA, ODMA, TDMA and SeDMA on serotonin 2A and 2B (5-HT$_{2A}$ and 5-HT$_{2B}$) receptors.** (**A**) At 5-HT$_{2A}$ receptor, the maximum activation values (as a % of 5-HT 1 µM activation) were: MDMA = 15.9%; ODMA=1.8%; TDMA=6.3%; SeDMA=6.1%. (**B**) At 5-HT$_{2B}$ receptor, the maximum activation values (as a % of 5-HT 1 µM activation) were: MDMA = 15.4%; ODMA = 2.3%; TDMA = 8.5%; SeDMA=7.9%. Data were normalized to the maximal fluorescence signal elicited by 5-HT 1 µM, 30 seconds after its application (max. value). Data are mean ± SEM (n=4).

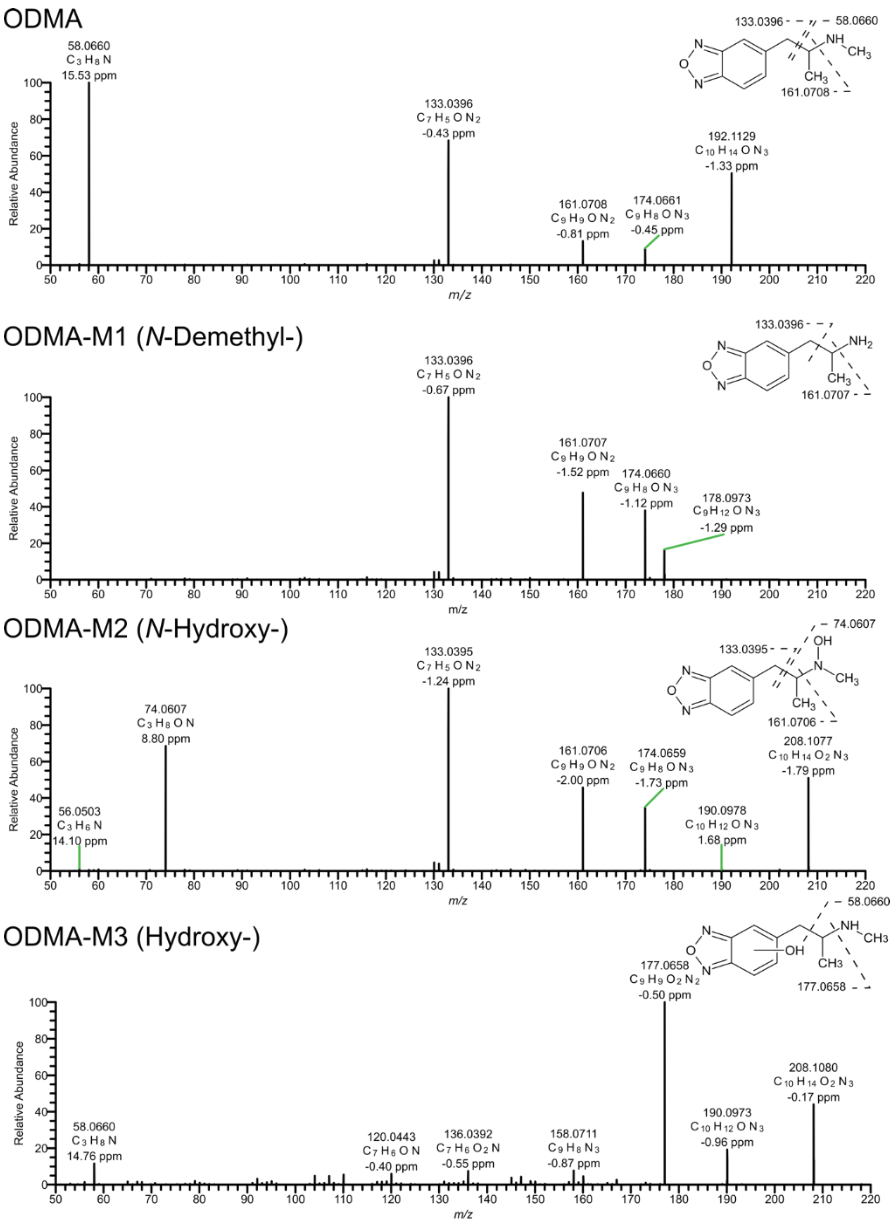

**Figure S5. LC-HRMS/MS spectra of ODMA and its metabolites identified in pooled human liver microsomes.** Metabolites are ordered by increasing mass. Metabolite-IDs correspond to Table 1. Fragments with accurate mass, calculated elemental formular, and mass error value in parts per million (ppm). *N*-demethylation is indicated by the absence of fragment ion (FI) at *m/z* 58.0660 ($C_3H_8N$) in comparison to the parent compound. *N*-hydroxylation was identified by the occurrence of the highly abundant fragment ion at *m/z* 74.0607 ($C_3H_8ON$). The remaining fragments revealed an unmodified ring system. Additionally, the retention time compared to parent compound supported the theory, since *N*-oxides are less hydrophilic and therefore, elute earlier using HILIC column whereases a hydroxylation leads usually to longer retention times. Monohydroxylation at the benzoxadiazole ring system was identified by the absence of FI at *m/z* 133.0396 ($C_7H_5ON_2$) compared to parent compound. Additionally, the occurrence of the FI at *m/z* 58.0660 ($C_3H_8N$), suggesting a hydroxylation on the ring system and not on the alkyl side chain.

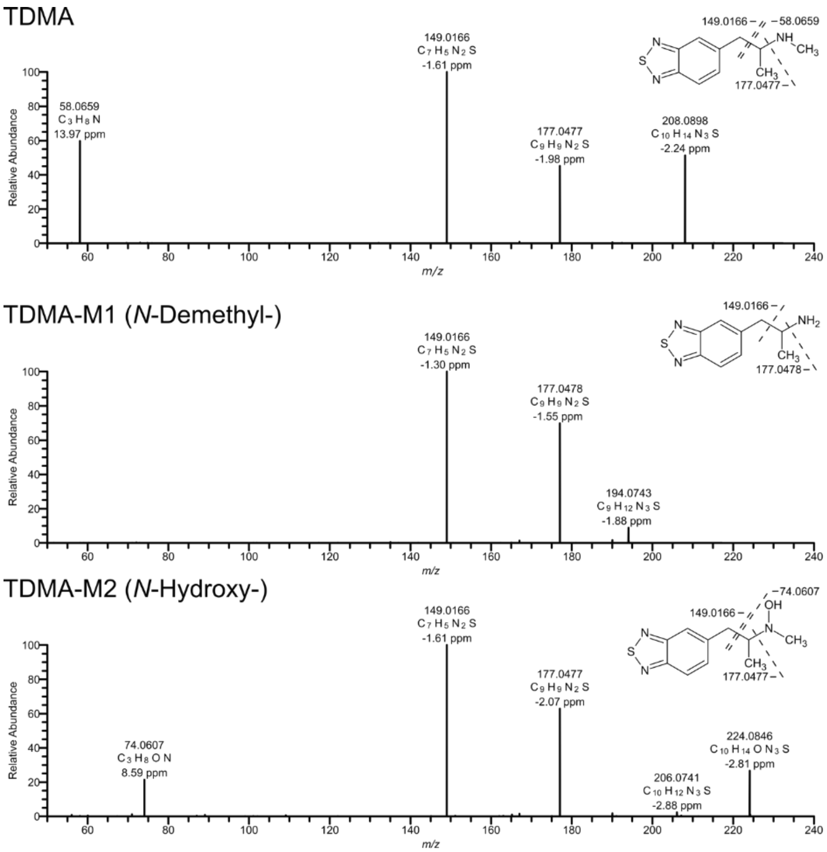

**Figure S6. LC-HRMS/MS spectra of TDMA and its metabolites identified in pooled human liver microsomes.** Metabolites are ordered by increasing mass. Metabolite-IDs correspond to Table 1. Fragments with accurate mass, calculated elemental formular, and mass error value in parts per million (ppm). Similar to ODMA in Figure S5, the lack of a fragment ion (FI) at *m/z* 58.0660 ($C_7H_5ON_2$) indicated the *N*-demethylation. *N*-Hydroxylation was also identified by the occurrence of FI at *m/z* 74.0607 ($C_3H_8ON$) along with the change in retention time as discussed above.

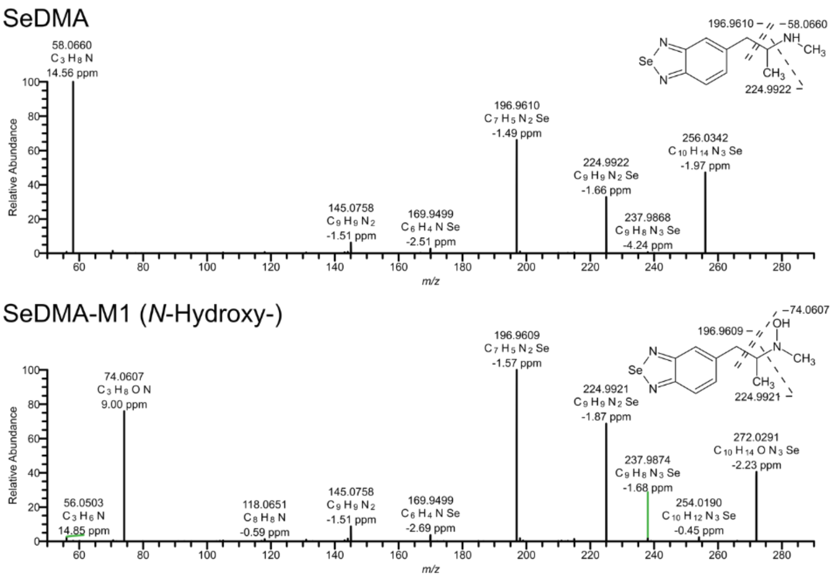

**Figure S7. LC-HRMS/MS spectra of SeDMA and its metabolite identified in pooled human liver microsomes.** Metabolites are ordered by increasing mass. Metabolite-IDs correspond to Table 1. Fragments with accurate mass, calculated elemental formular, and mass error value in parts per million (ppm). Identification of the metabolic reaction followed the same principles as for ODMA and TDMA (Figure S5 and S6).

### 3. Electron ionization mass spectra and NMR spectra

#### 3.1. Electron ionization mass spectra

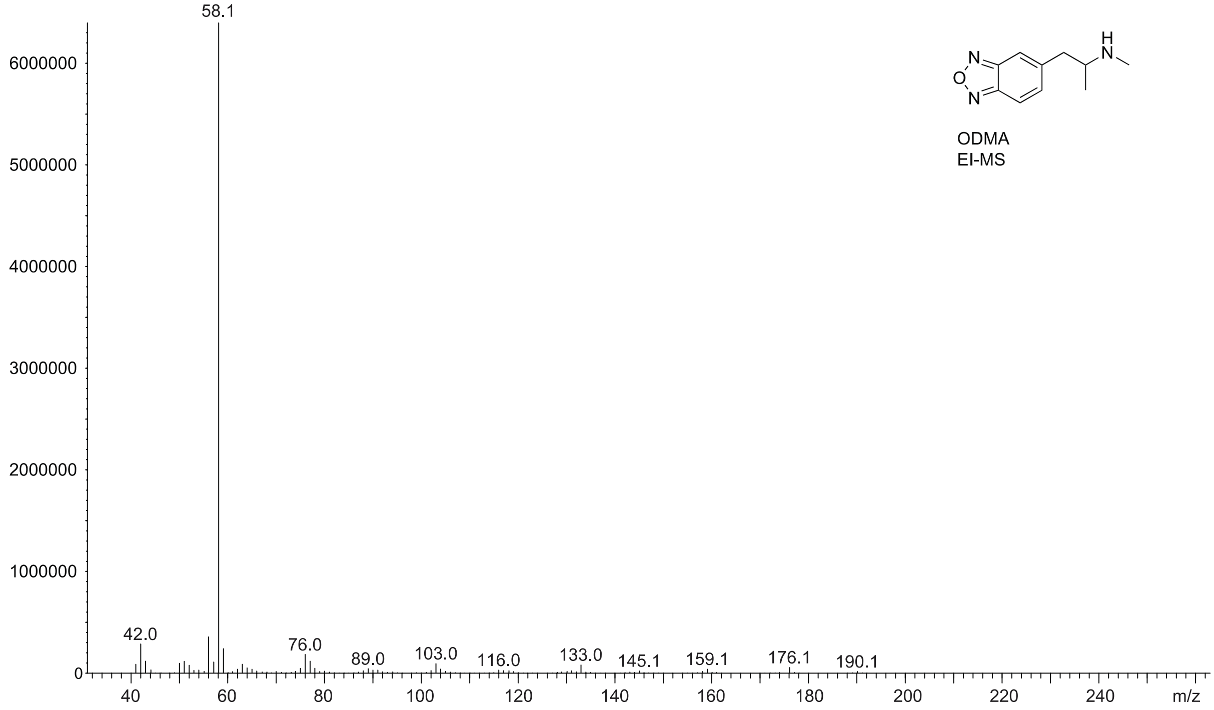

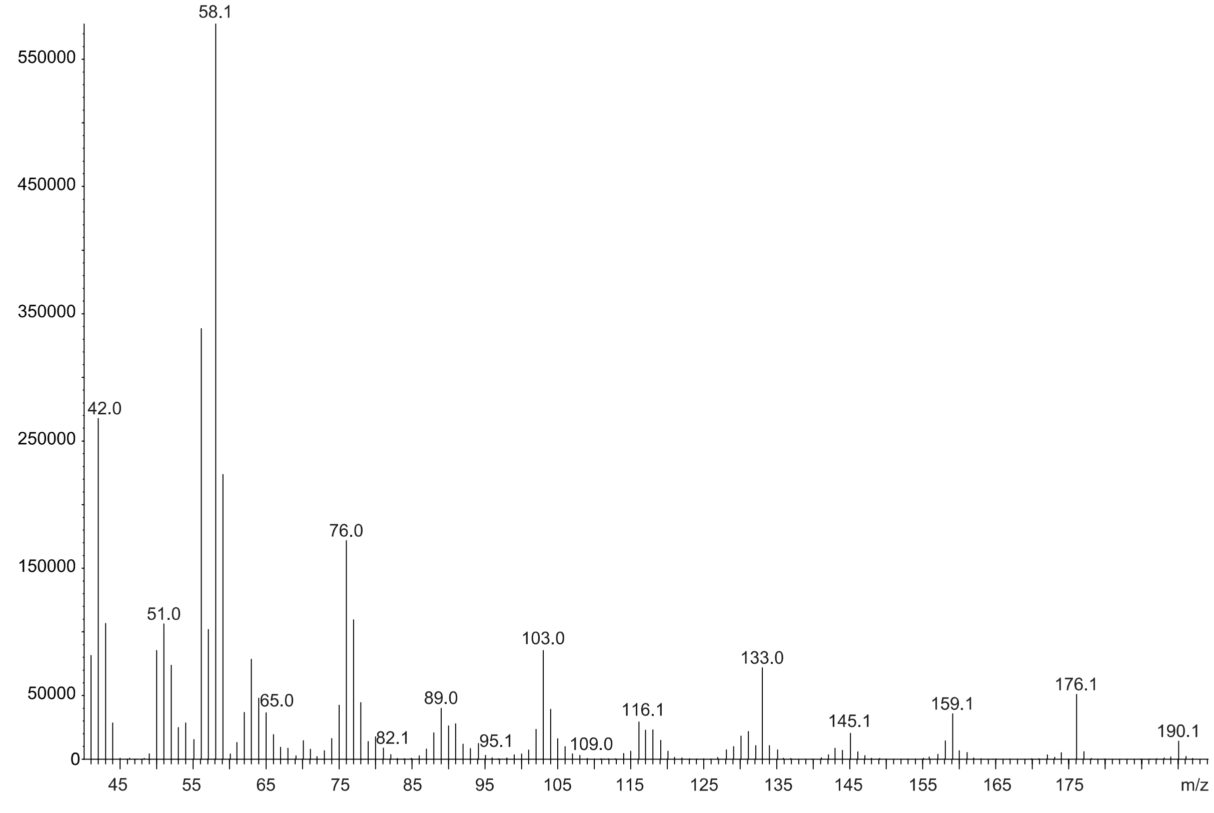

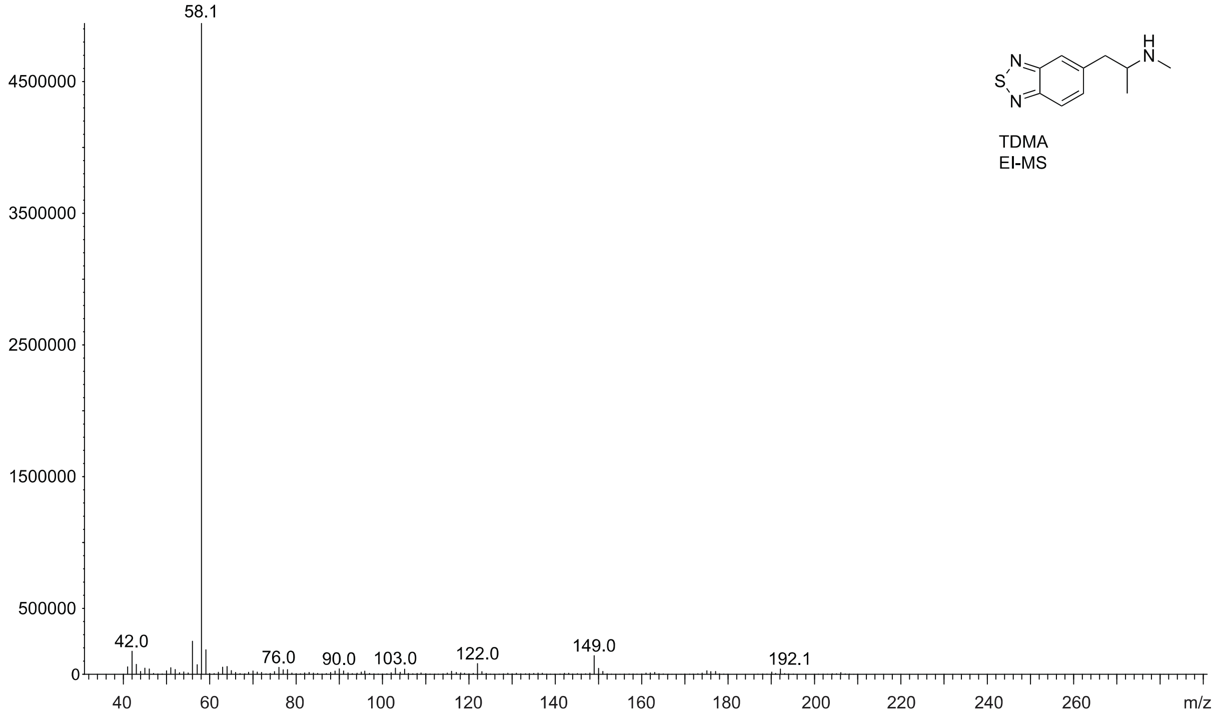
58.1
4500000
3500000
2500000
1500000
500000
0
42.0
76.0
90.0
103.0
122.0
149.0
192.1
40
60
80
100
120
140
160
180
200
220
240
260
m/z
TDMA
EI-MS

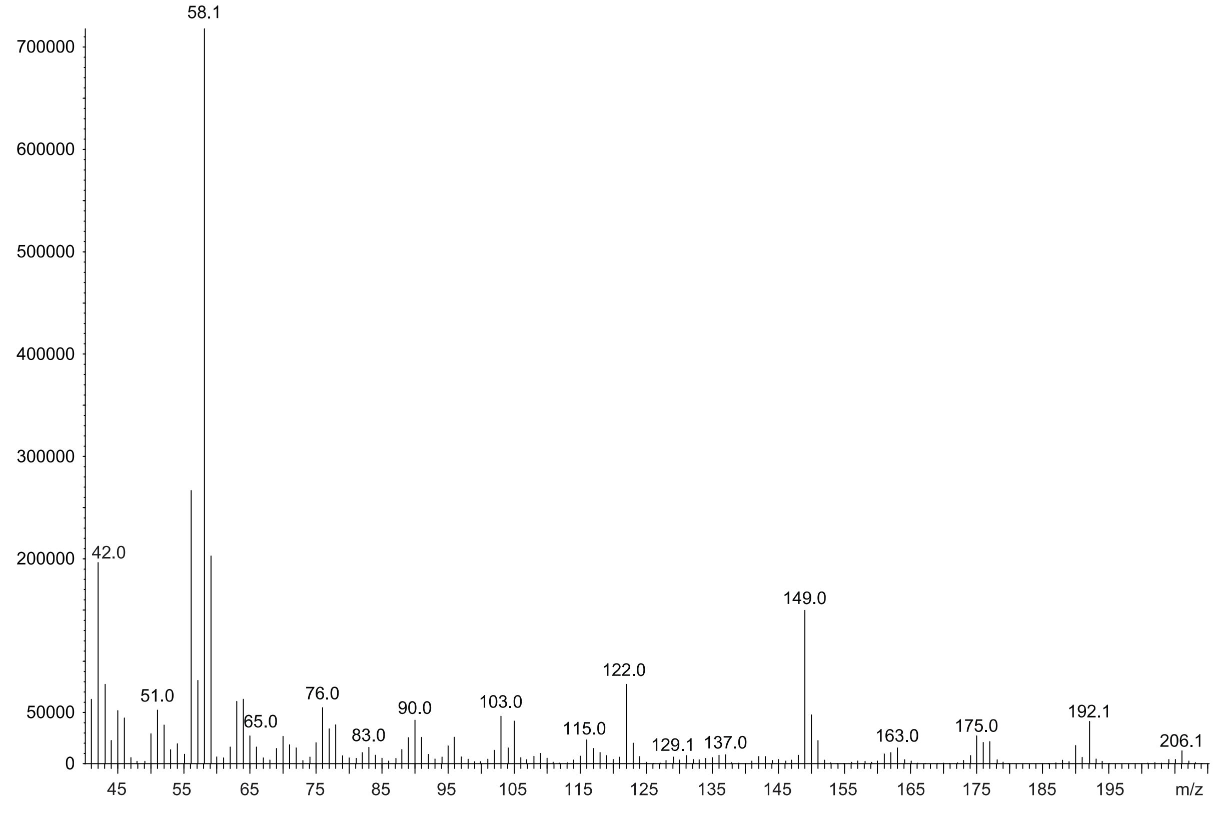
58.1
700000
600000
500000
400000
300000
200000
50000
0
42.0
51.0
65.0
76.0
83.0
90.0
103.0
115.0
122.0
129.1
137.0
149.0
163.0
175.0
192.1
206.1
45
55
65
75
85
95
105
115
125
135
145
155
165
175
185
195
m/z

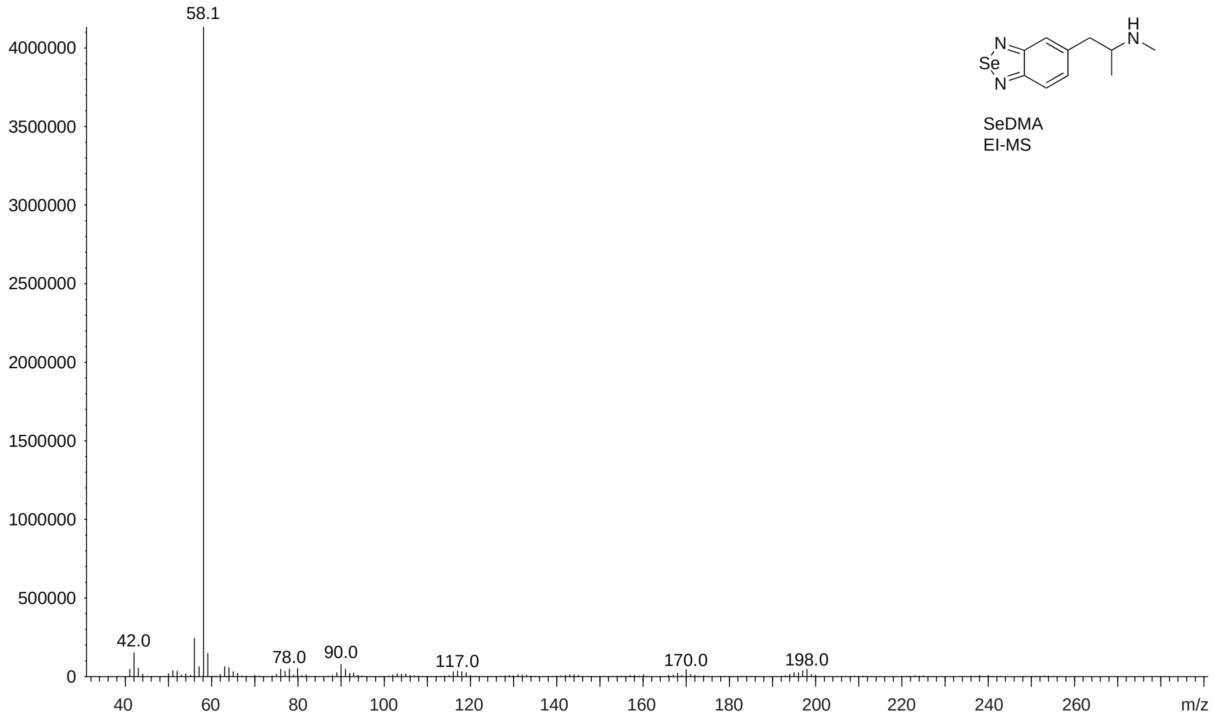
58.1
4000000
3500000
3000000
2500000
2000000
1500000
1000000
500000
0
42.0
78.0
90.0
117.0
170.0
198.0
40
60
80
100
120
140
160
180
200
220
240
260
m/z
H
N
N
Se
N
SeDMA
EI-MS

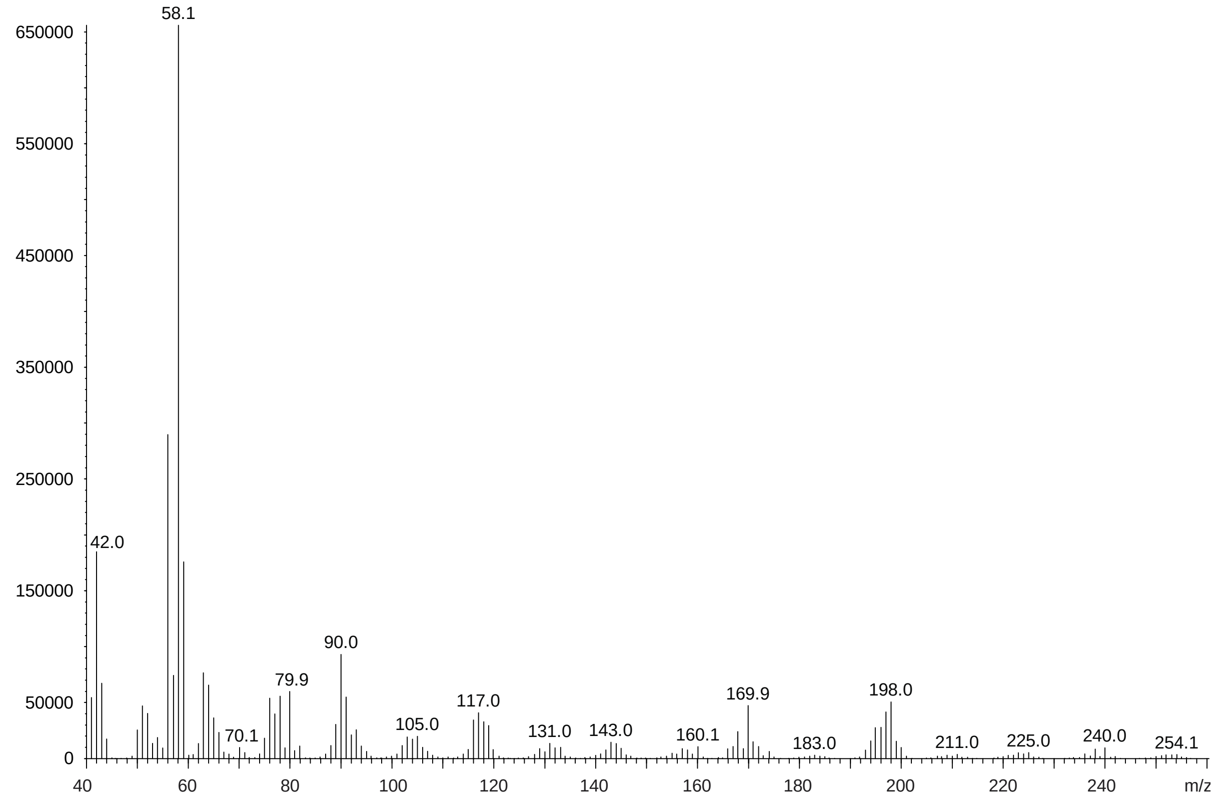
58.1
650000
550000
450000
350000
250000
150000
50000
0
42.0
70.1
79.9
90.0
105.0
117.0
131.0
143.0
160.1
169.9
183.0
198.0
211.0
225.0
240.0
254.1
40
60
80
100
120
140
160
180
200
220
240
m/z

##### 3.2. NMR Spectra

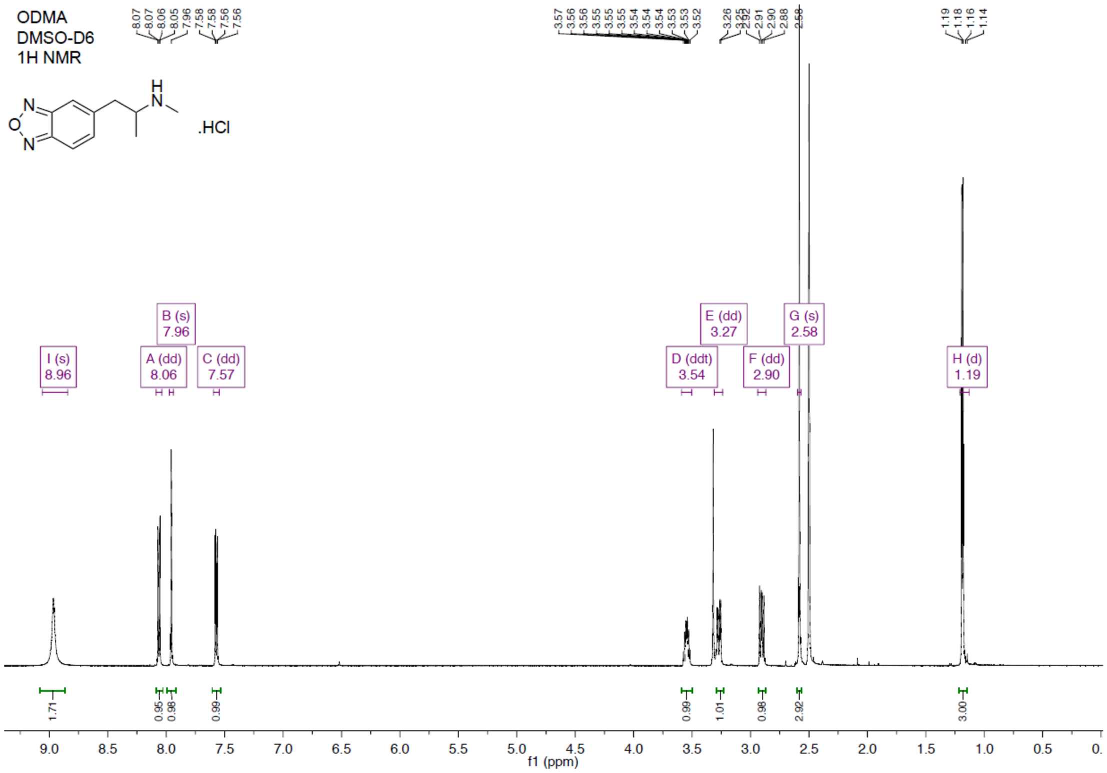

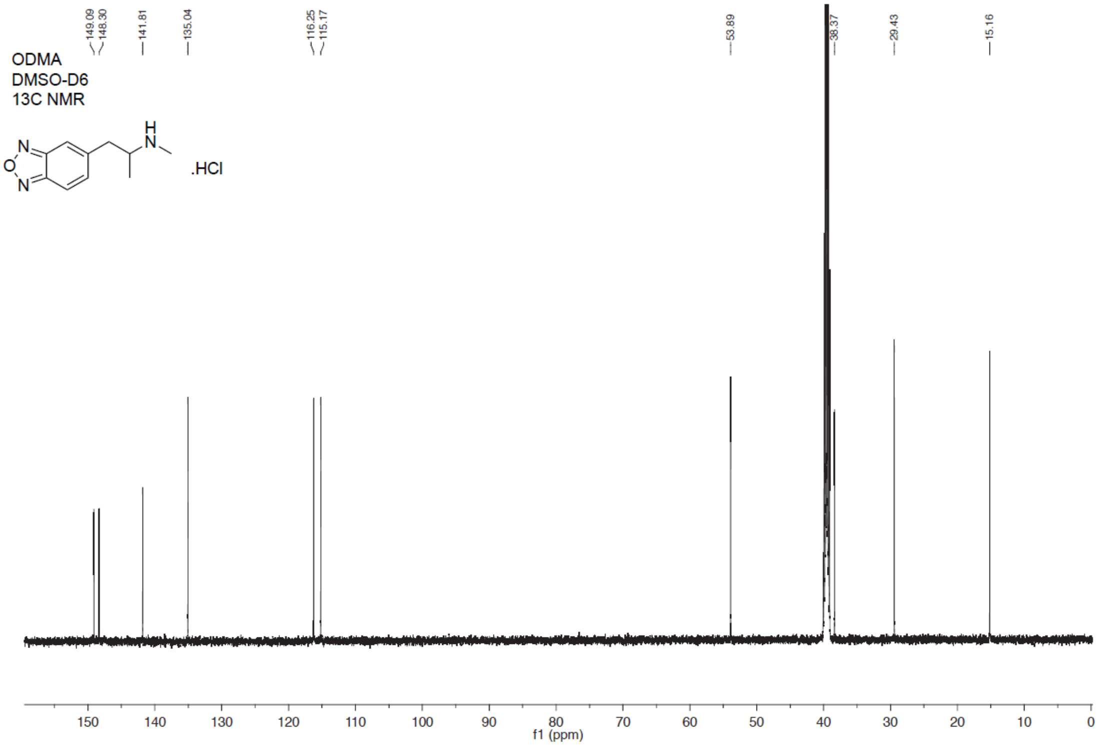

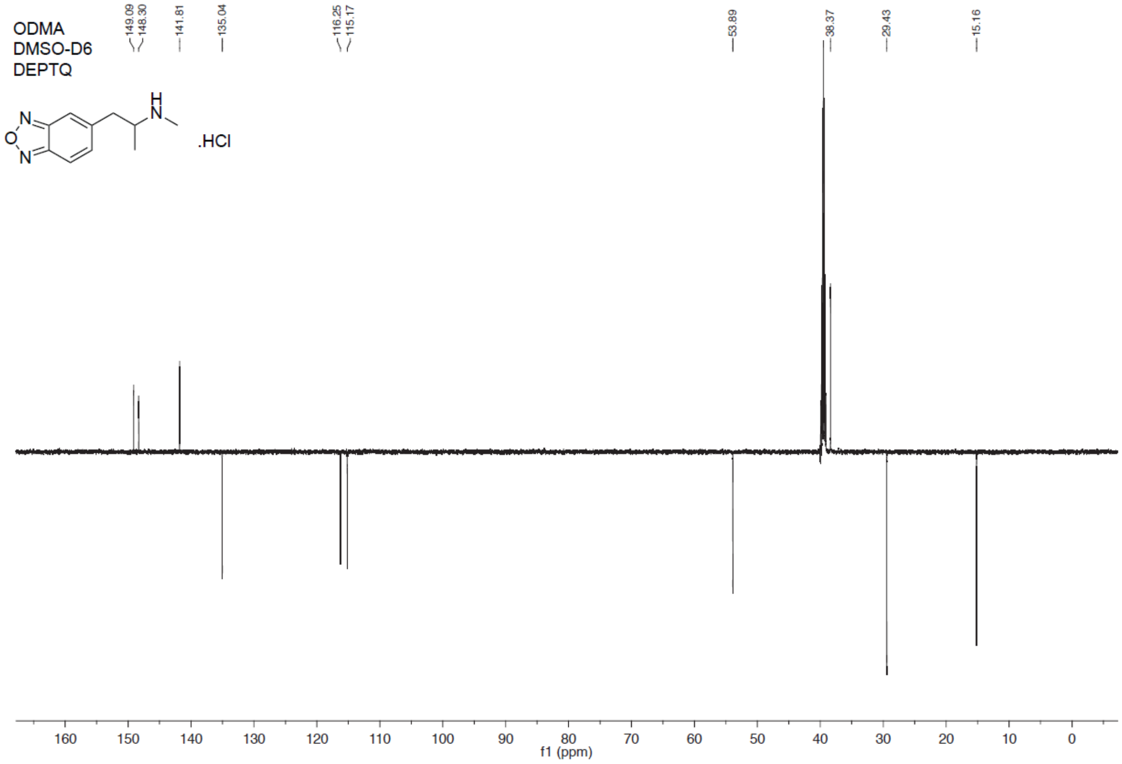
ODMA
DMSO-D6
DEPTQ
.HCl
149.09
148.30
141.81
135.04
116.25
115.17
53.89
38.37
29.43
15.16
f1 (ppm)

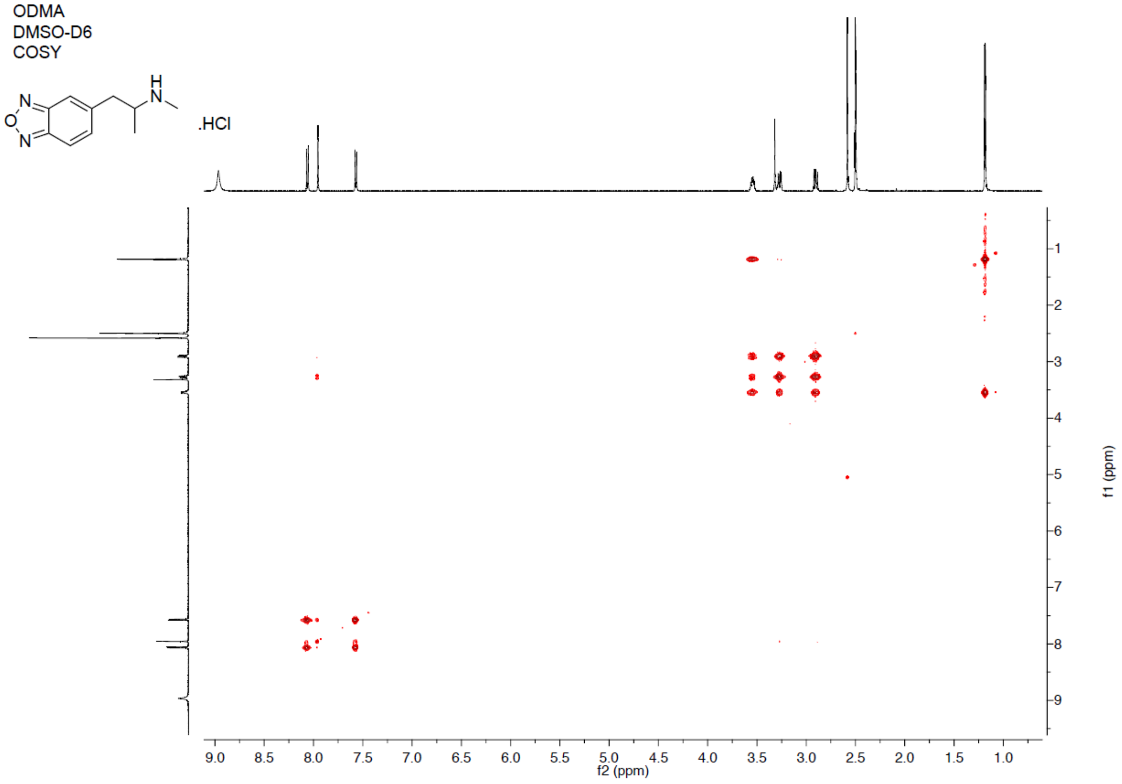
ODMA
DMSO-D6
COSY
.HCl
f2 (ppm)
f1 (ppm)

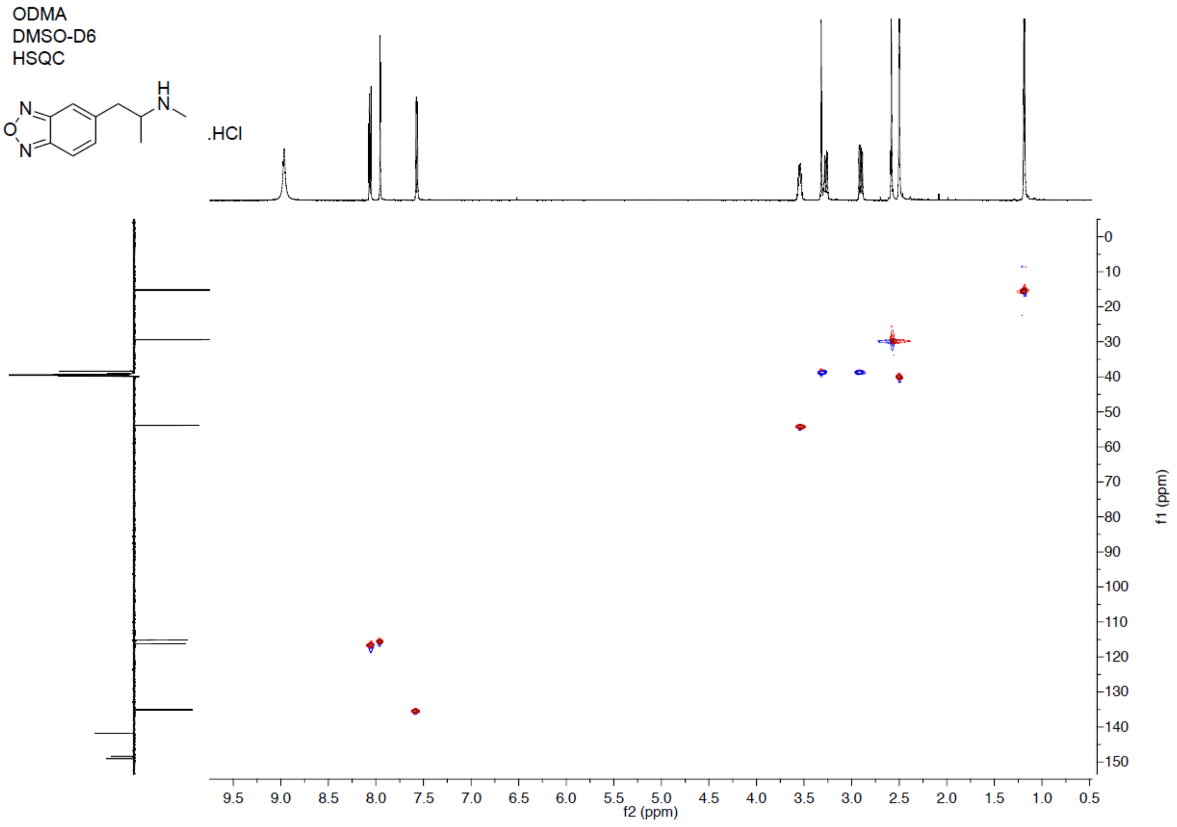
ODMA
DMSO-D6
HSQC
.HCl
f2 (ppm)
f1 (ppm)

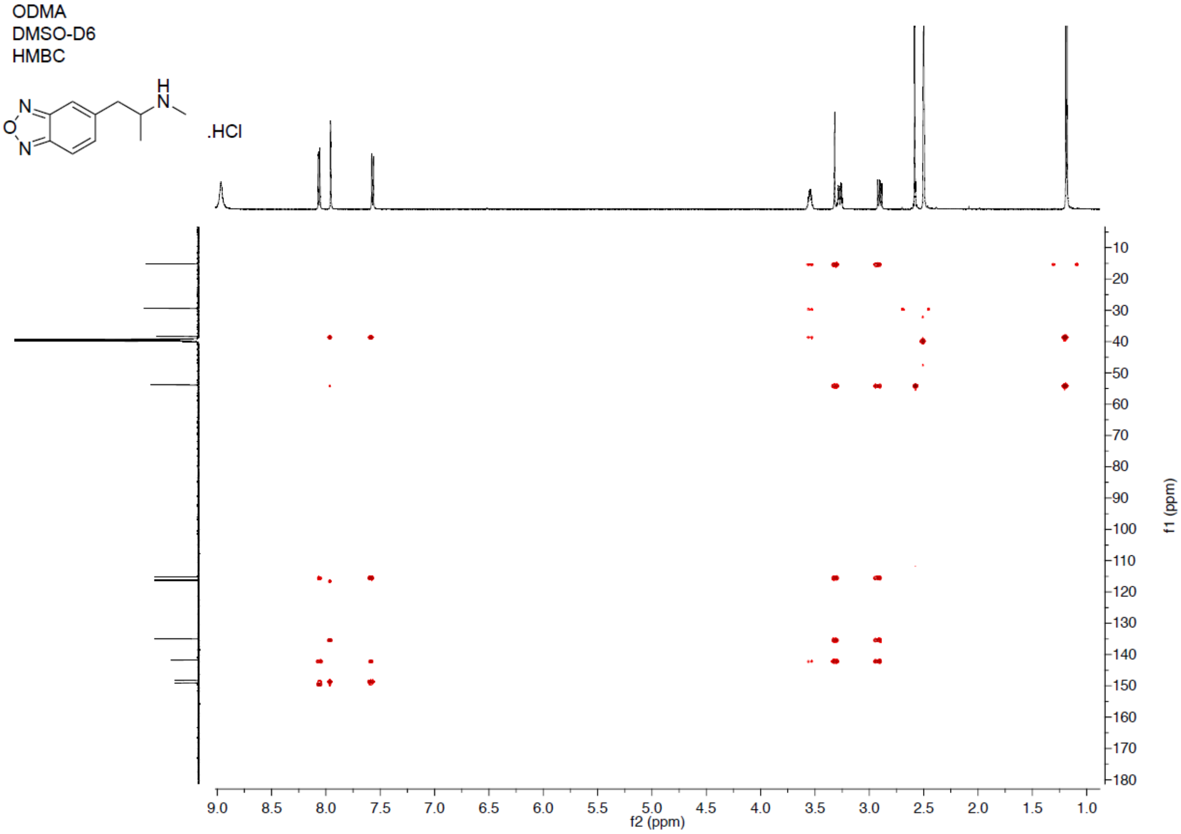
ODMA
DMSO-D6
HMBC
.HCl
f2 (ppm)
f1 (ppm)

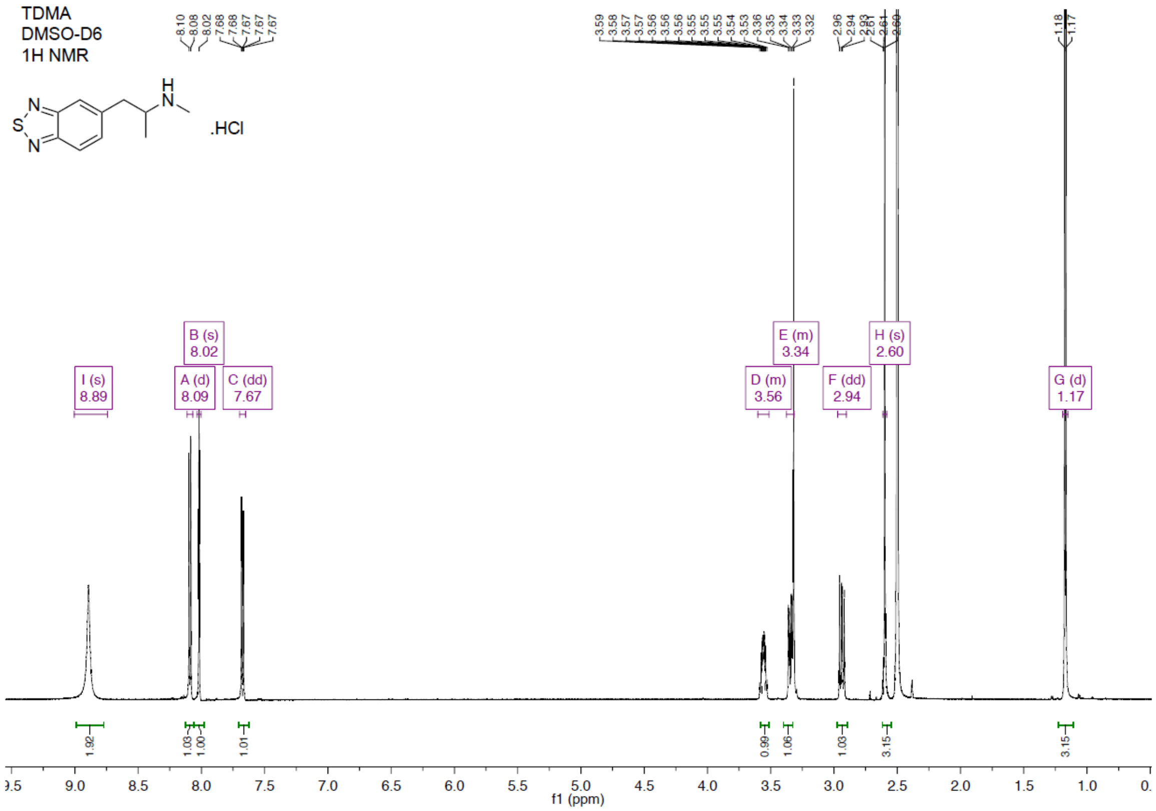
TDMA
DMSO-D6
1H NMR
.HCl
I (s) 8.89
B (s) 8.02
A (d) 8.09
C (dd) 7.67
D (m) 3.56
E (m) 3.34
F (dd) 2.94
H (s) 2.60
G (d) 1.17
f1 (ppm)

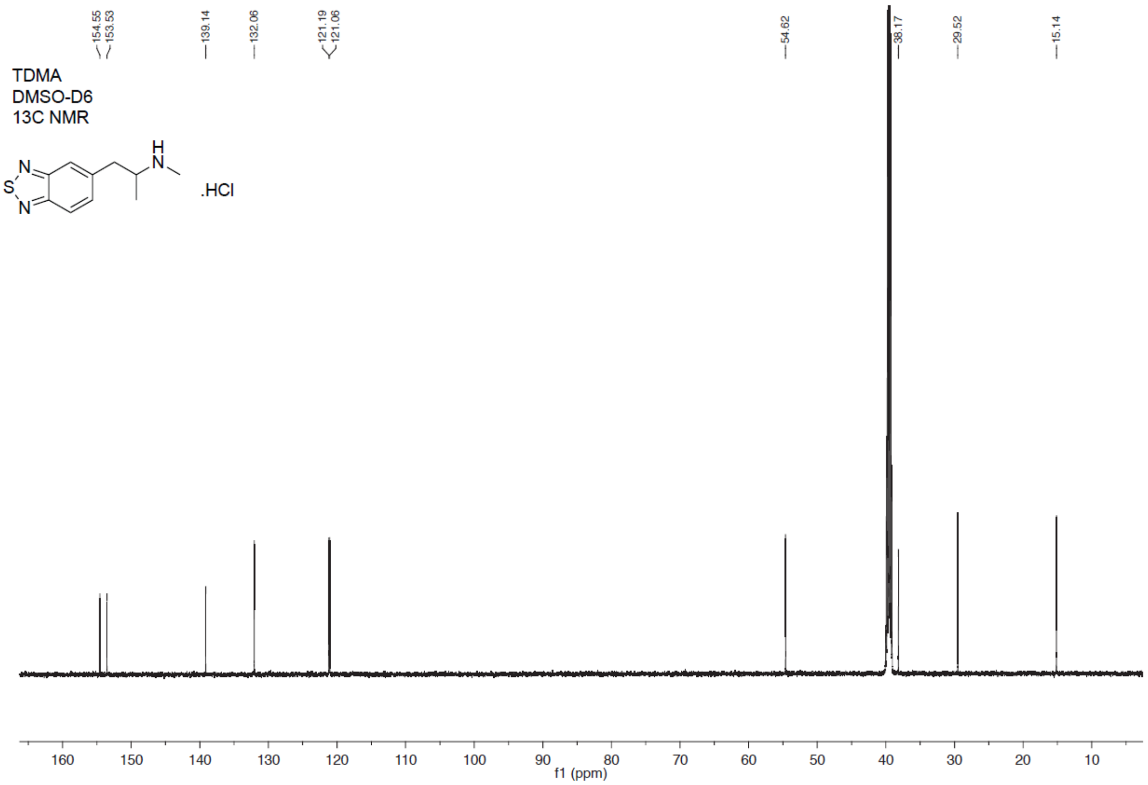
154.55
153.53
139.14
132.06
121.19
121.06
54.62
38.17
29.52
15.14
TDMA
DMSO-D6
13C NMR
.HCl
f1 (ppm)

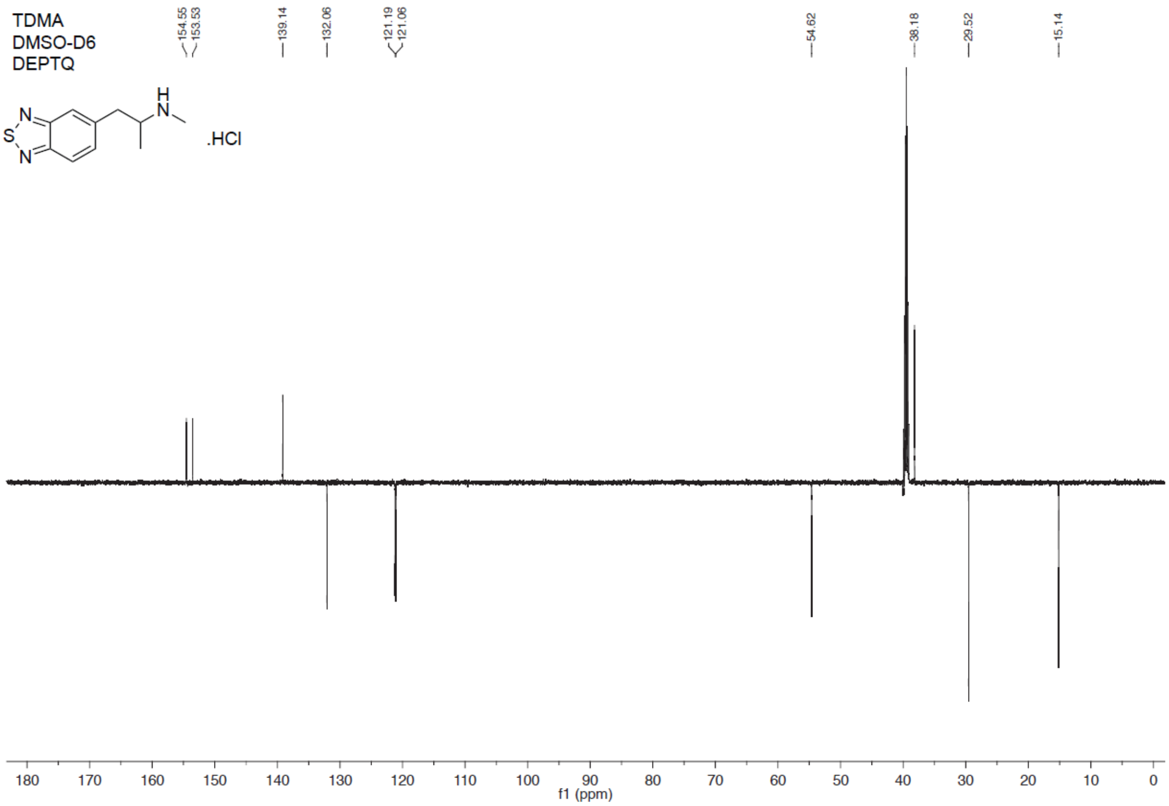
TDMA
DMSO-D6
DEPTQ
.HCl
154.55
153.53
139.14
132.06
121.19
121.06
54.62
38.18
29.52
15.14
f1 (ppm)

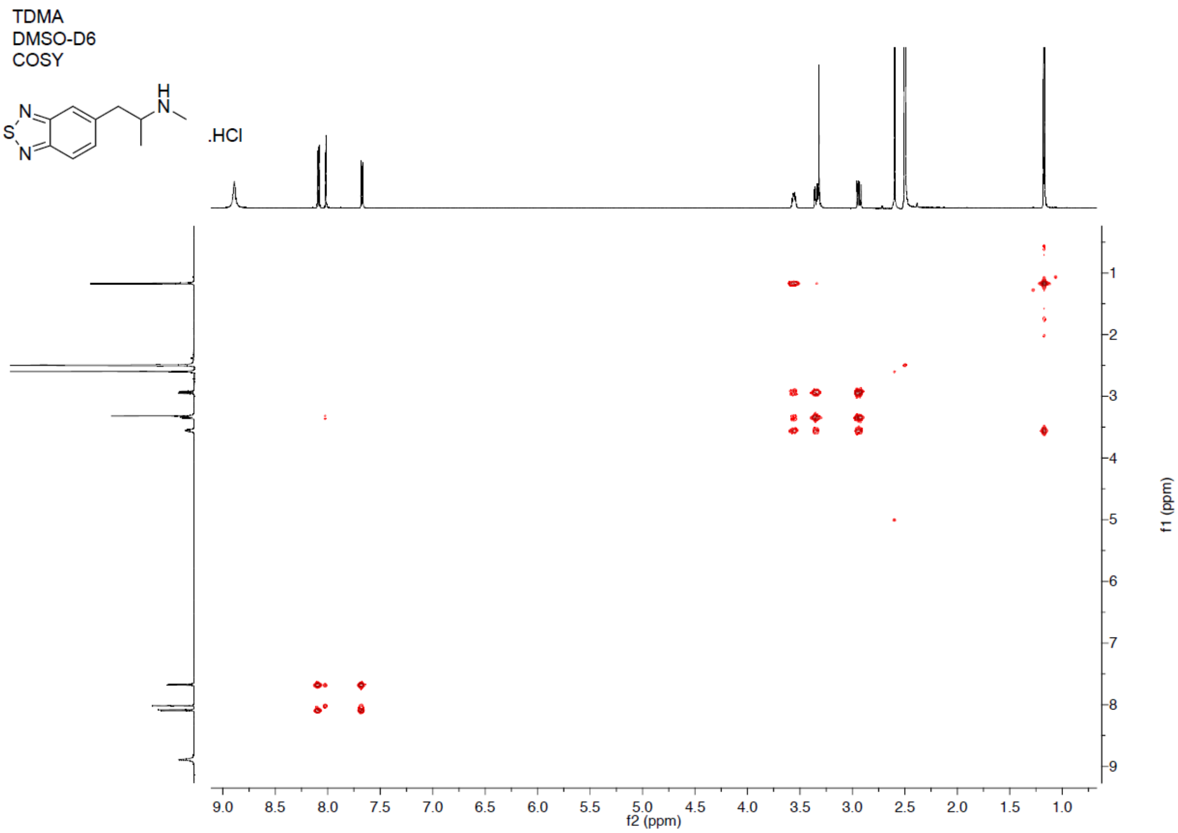
TDMA
DMSO-D6
COSY
.HCl
f2 (ppm)
f1 (ppm)

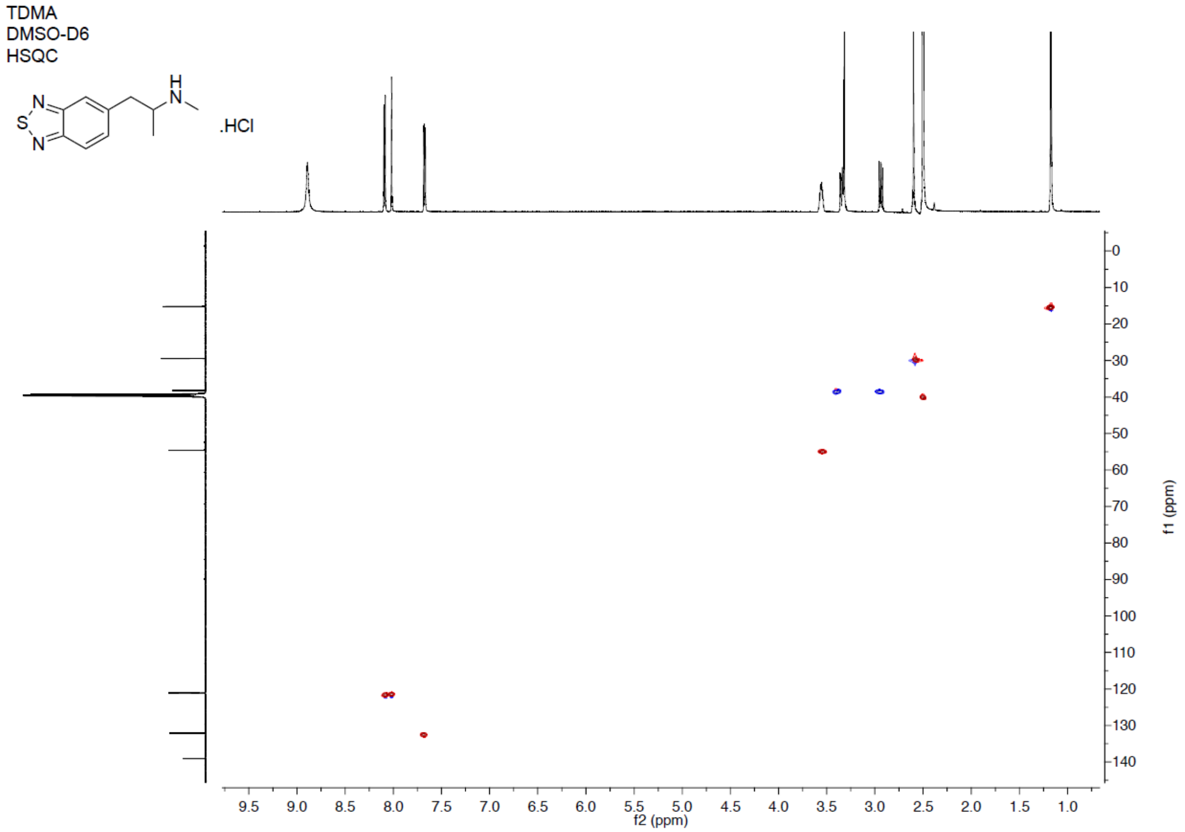
TDMA
DMSO-D6
HSQC
.HCl
f2 (ppm)
f1 (ppm)

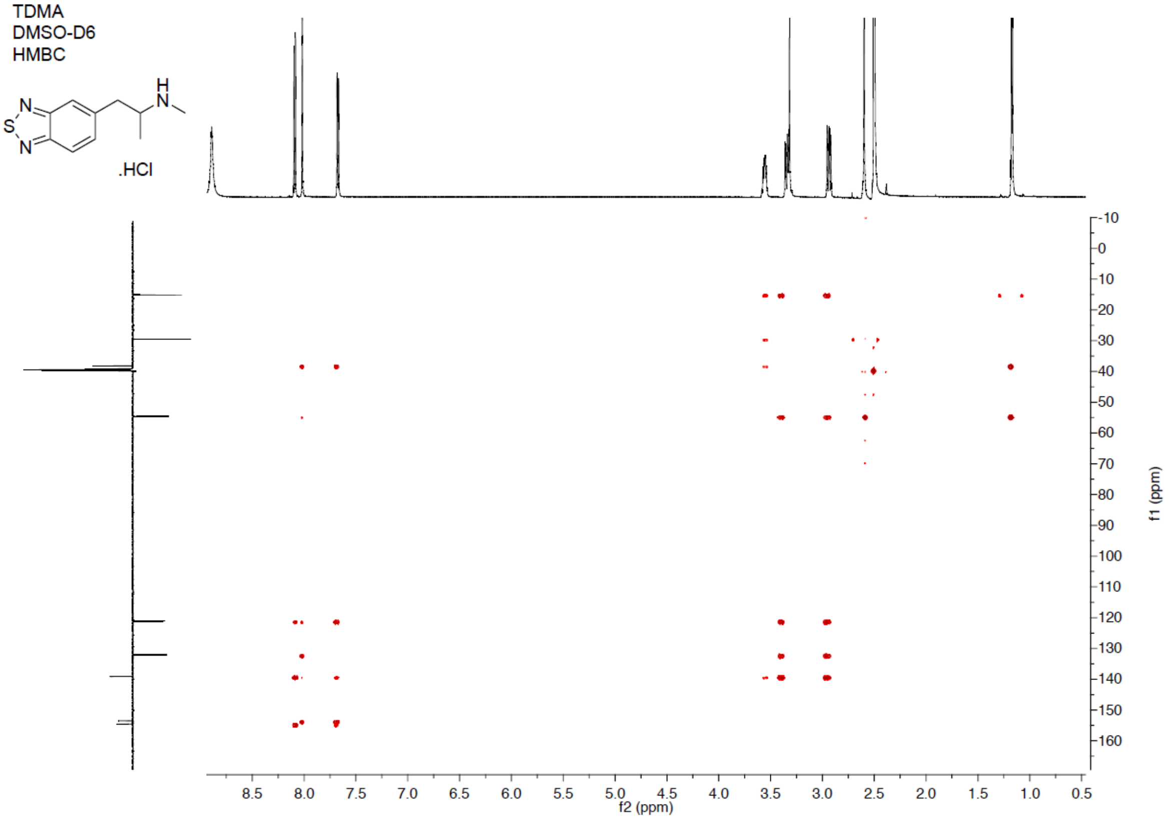
TDMA
DMSO-D6
HMBC
.HCl
f2 (ppm)
f1 (ppm)

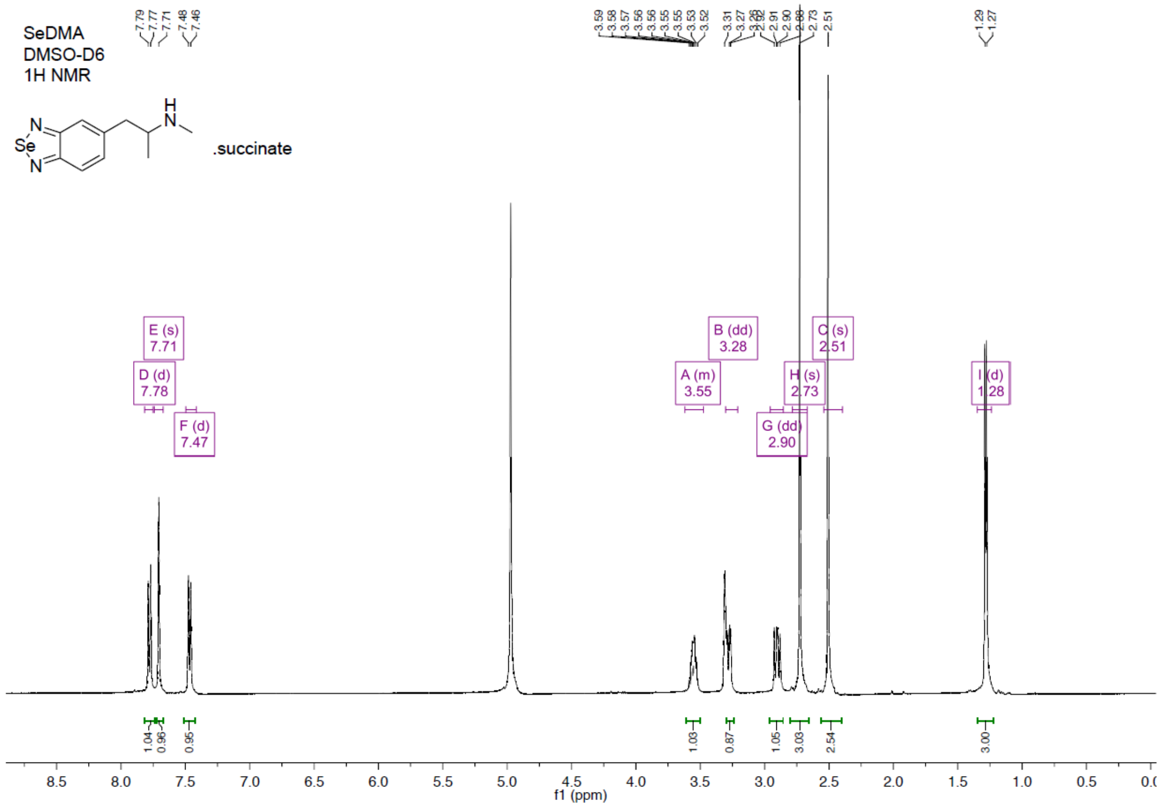
SeDMA
DMSO-D6
1H NMR
.succinate
E (s) 7.71
D (d) 7.78
F (d) 7.47
B (dd) 3.28
A (m) 3.55
C (s) 2.51
H (s) 2.73
G (dd) 2.90
I (d) 1.28
f1 (ppm)

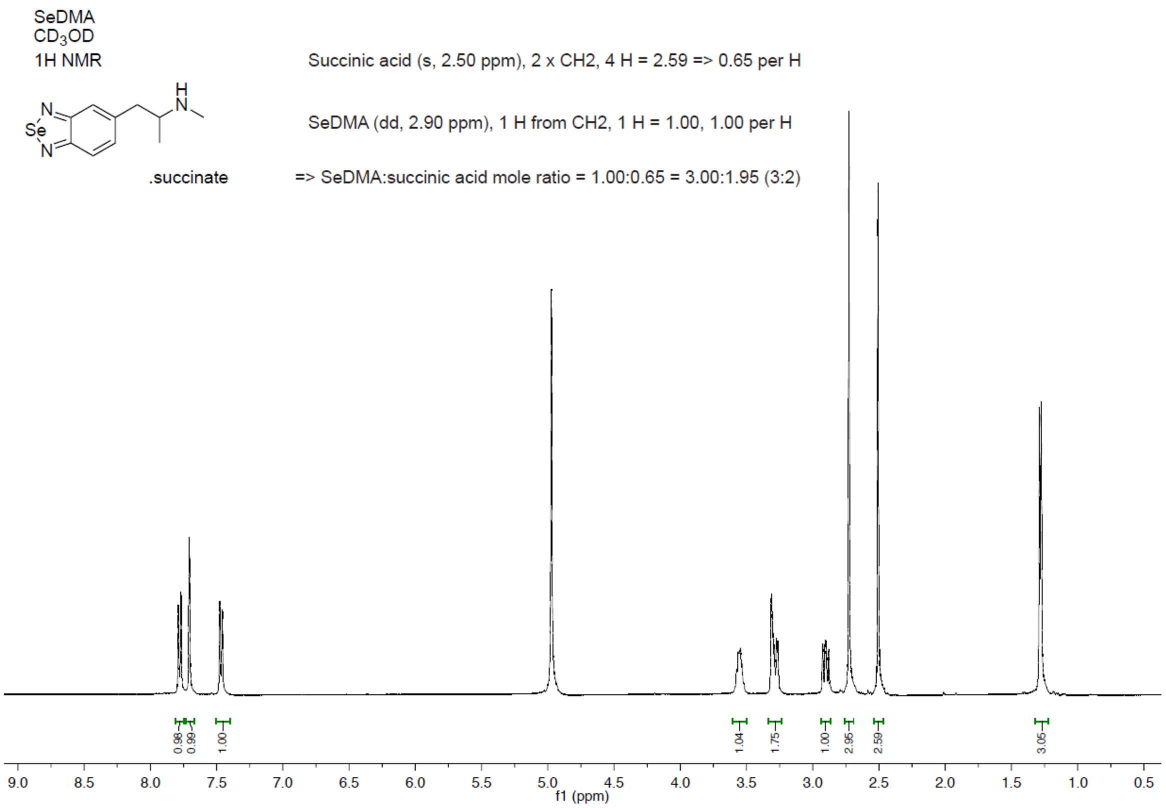
SeDMA
$CD_3OD$
1H NMR
.succinate
Succinic acid (s, 2.50 ppm), 2 x CH2, 4 H = 2.59 => 0.65 per H
SeDMA (dd, 2.90 ppm), 1 H from CH2, 1 H = 1.00, 1.00 per H
=> SeDMA:succinic acid mole ratio = 1.00:0.65 = 3.00:1.95 (3:2)
f1 (ppm)

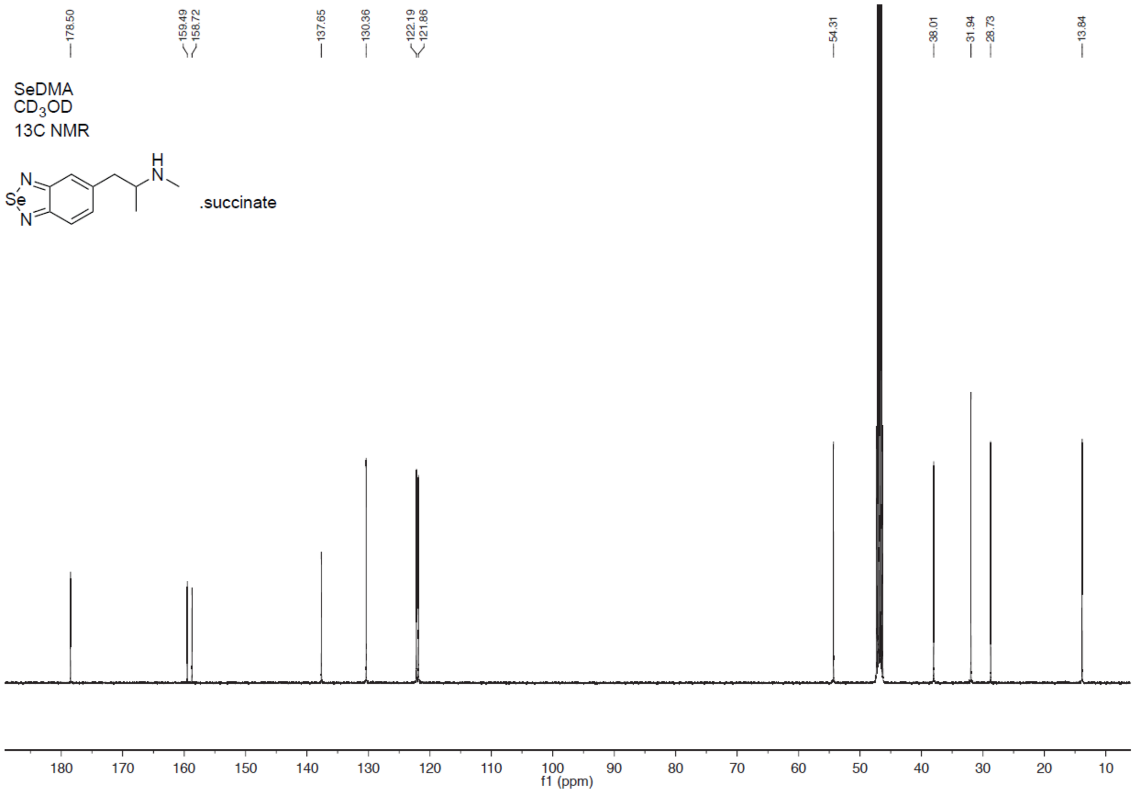
178.50
159.49
158.72
137.65
130.36
122.19
121.86
54.31
38.01
31.94
28.73
13.84
SeDMA
CD3OD
13C NMR
.succinate
f1 (ppm)

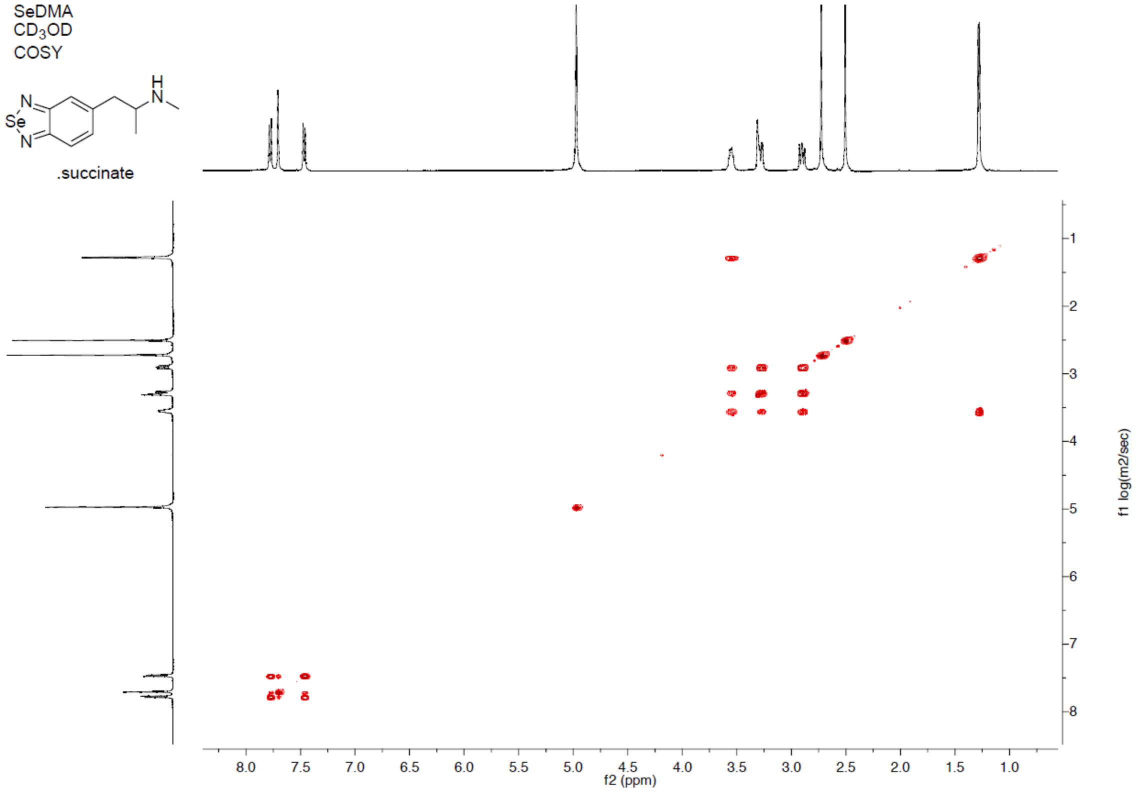
SeDMA
CD3OD
COSY
.succinate
f2 (ppm)
f1 log(m2/sec)

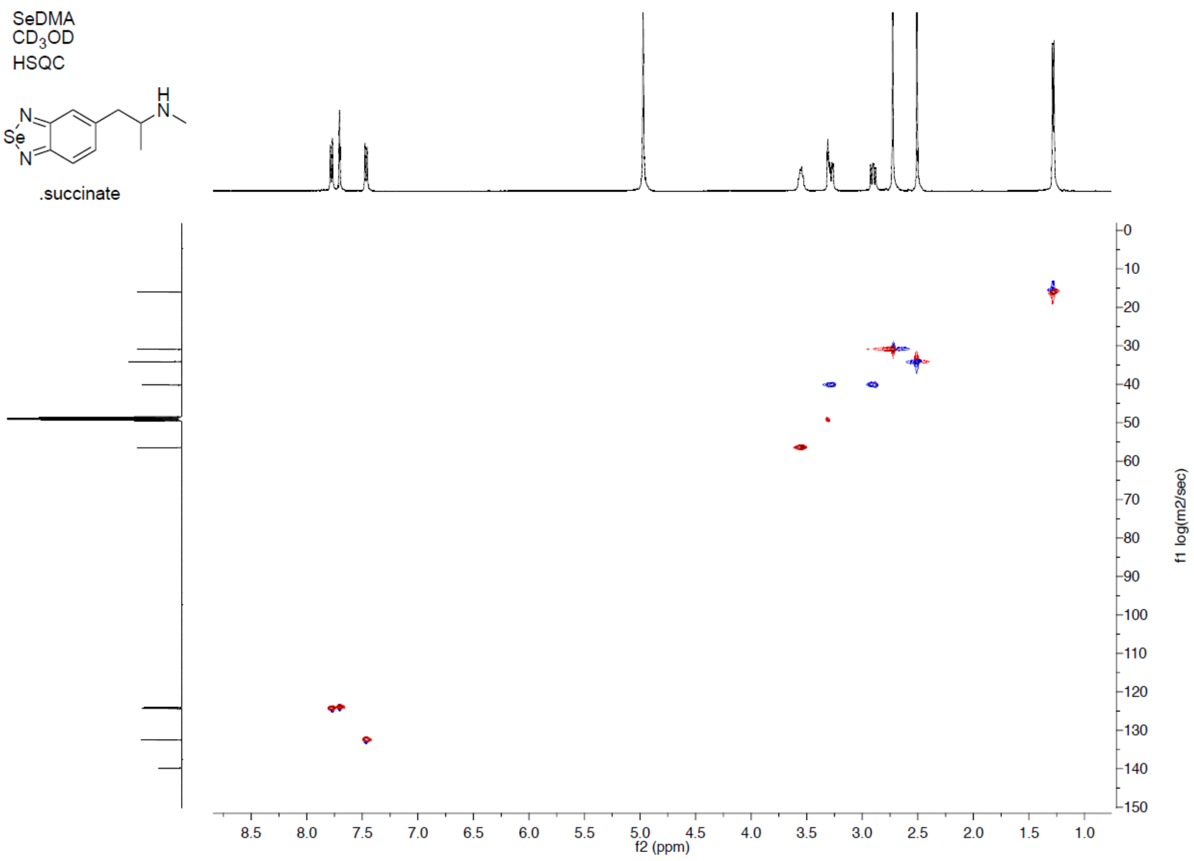
SeDMA
CD3OD
HSQC
.succinate
f2 (ppm)
f1 log(m2/sec)

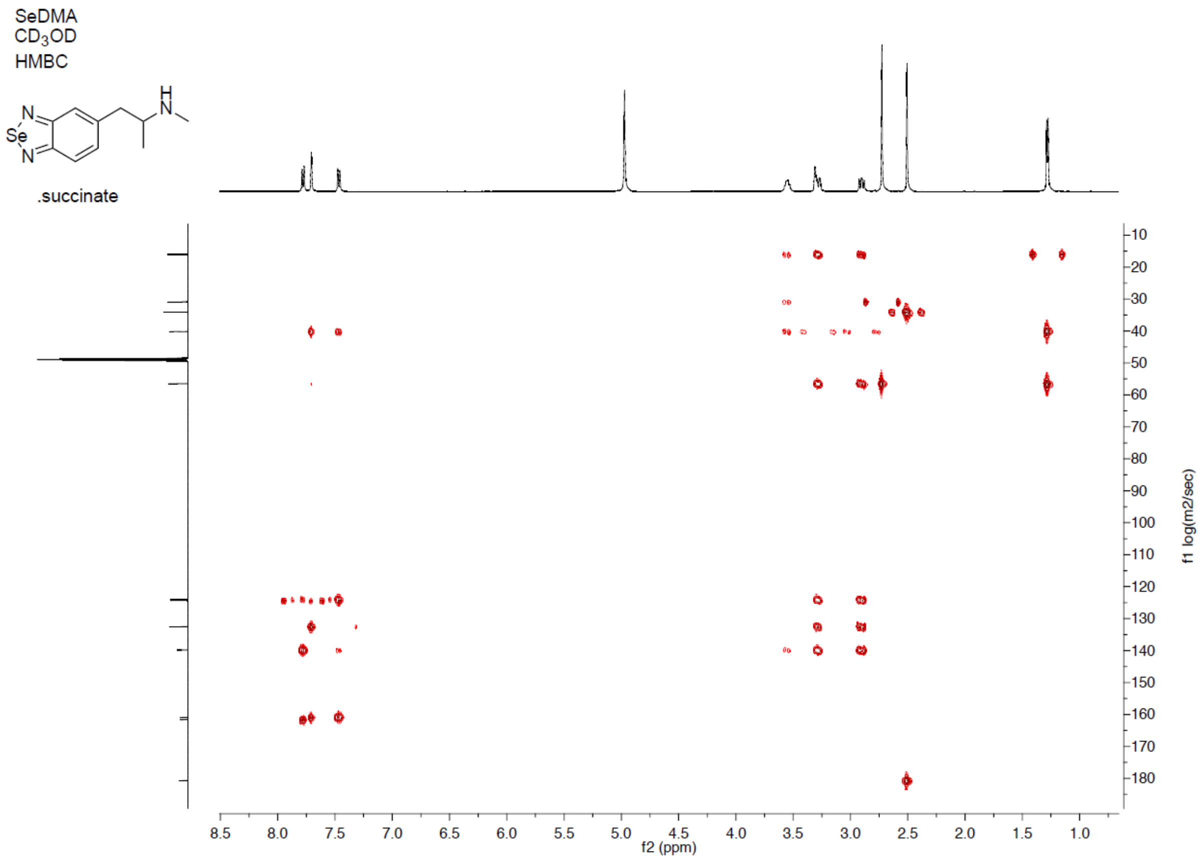
SeDMA
CD3OD
HMBC
.succinate
f2 (ppm)
f1 log(m2/sec)
